# An injury-induced tissue niche shaped by mesenchymal plasticity coordinates the regenerative and disease response in the lung

**DOI:** 10.1101/2024.02.26.582147

**Authors:** Dakota L. Jones, Michael P. Morley, Xinyuan Li, Yun Ying, Fabian L. Cardenas-Diaz, Shanru Li, Su Zhou, Sarah E. Schaefer, Ullas V. Chembazhi, Ana Nottingham, Susan Lin, Edward Cantu, Joshua M. Diamond, Maria C. Basil, Andrew E. Vaughan, Edward E. Morrisey

## Abstract

Severe lung injury causes basal stem cells to migrate and outcompete alveolar stem cells resulting in dysplastic repair and a loss of gas exchange function. This “stem cell collision” is part of a multistep process that is now revealed to generate an <u>i</u>njury-induced <u>t</u>issue ni<u>ch</u>e (iTCH) containing Keratin 5+ epithelial cells and plastic Pdgfra+ mesenchymal cells. Temporal and spatial single cell analysis reveals that iTCHs are governed by mesenchymal proliferation and Notch signaling, which suppresses Wnt and Fgf signaling in iTCHs. Conversely, loss of Notch in iTCHs rewires alveolar signaling patterns to promote euplastic regeneration and gas exchange. The signaling patterns of iTCHs can differentially phenotype fibrotic from degenerative human lung diseases, through apposing flows of FGF and WNT signaling. These data reveal the emergence of an injury and disease associated iTCH in the lung and the ability of using iTCH specific signaling patterns to discriminate human lung disease phenotypes.

## Introduction

The lung is generally quiescent during normal life but upon severe injury or in disease states, specific cell types in the lung alveolus re-enter the cell cycle, differentiate, and replenish lost cells to re-establish respiratory function. How this process occurs and the cellular constituents that are involved remain unclear. Disruption of this multi-step regenerative process can lead to permanent loss of respiratory function and chronic lung diseases.

Respiratory viral infections, such as influenza A, are a leading global cause of infectious morbidity and mortality(*1*, *2*). Respiratory illness caused by influenza follows a stereotypical course with infection of lung epithelium, including alveolar type 2 (AT2) cells, cytotoxic destruction of alveolar tissue, dramatic inflammatory infiltration, and early immune-responsive fibroblast activation(*3*). This tissue destruction is followed by proliferation of multiple alveolar cell types including AT2 cells, cellular differentiation including AT2 cells differentiating into alveolar type 1 (AT1) cells, and re-establishment of normal alveolar architecture. In mild to moderate lung injury, most of the destroyed alveolar tissue can be regenerated through what has been referred to as the euplastic regenerative response, resulting in the re-establishment of alveolar structure and restoration of gas-exchange function(*4*, *5*). However, following severe injury, a dysplastic response can also occur which entails the egression of rare intrapulmonary basal stem cells out of the airways into the alveolar parenchyma. The dysplastic basal cells migrate, proliferate, and rapidly expand into regions of severely damaged alveolar regions of the lung. These dysplastic epithelial stem cells express Keratin 5 (Krt5) and form regions referred to as “pods”, which persist and can be found in patients who survived severe influenza infections(*6*–*8*). Furthermore, Krt5+ dysplastic epithelium in the alveolar compartment is also found in some chronic lung diseases including COVID-19-induced fibrosis(*8*). Understanding how the lung balances euplastic versus dysplastic tissue reactions is central to promoting improved lung function post-injury as well as developing new therapies for chronic lung diseases.

The contribution of non-epithelial cells in balancing euplastic versus dysplastic tissue responses to acute lung injury remains unclear. Lineage tracing shows the emergence of rare Trp63+ intrapulmonary basal cells to generate the dysplastic epithelial response post-viral injury, resulting in these cells exiting their airway niche and migrating into the alveolar niche, which is occupied by completely different cell lineages and signaling networks(*7–9*). How the interactions between the alveolar and airway cell types within these two niches facilitate the emergence of this keratinized dysplasia remains unknown.

Here, we show that the dysplastic epithelial response after acute injury to the lung leads to the formation of a multi-cellular <u>i</u>njury-induced <u>t</u>issue ni<u>ch</u>e (iTCH). iTCHs are comprised of both Krt5+ dysplastic epithelium and alveolar Pdgfrb+ mesenchymal cells (also called alveolar fibroblast 2 or AF2), which are derived from alveolar Pdgfra+ cells (also called alveolar fibroblast 1 or AF1). This mesenchymal plasticity is unidirectional, with Pdgfra+ AF1 cells proliferating and differentiating into Pdgfrb+ AF2 cells, while Pdgfrb+ AF2 cells do not proliferate or generate Pdgfra+ AF1 cells. Pdgfra-derived Pdgfrb+ cells closely integrate into iTCHs and form a unique Notch mediated niche with the Krt5+ dysplastic epithelium. Suppression of cell proliferation or Notch signaling in Pdgfra+ cells prior to injury results in rewiring of the cell-cell communication axis in the alveolus, leading to a loss of iTCHs with a commensurate increase in Wnt and Fgf signaling flow resulting in increased alveolar euplastic regeneration. Using newly generated and existing multi-disease single cell transcriptional datasets, we show that iTCHs and NOTCH mediated dysplastic cell-cell communication pathways are observed in human fibrotic diseases including post-COVID-19 fibrosis and bleomycin induced human lung injury. In contrast, emphysematous degenerative lung diseases such as chronic obstructive pulmonary disease (COPD) and alpha-1 anti-trypsin (AAT) disease lack iTCHs and exhibit a reversal of cell-cell communication flows, demonstrating that a euplastic regenerative response in the alveolar niche dominates. These studies delineate an emergent niche that arises after injury and in specific human lung diseases, that dictates whether the tissue engages a dysplastic repair or euplastic regeneration response.

## Results

### Pdgfra expressing cells are the primary reactive mesenchyme after lung injury and exhibit progenitor cell function

While previous work has shown that the lung mesenchyme reacts to acute injury(*10*, *11*), its contribution towards alveolar regeneration is poorly understood. We analyzed our previously described mouse scRNA-seq data from adult animals and combined this with additional analysis of spatial localization of mesenchymal subsets(*12*). This analysis revealed two distinct populations of alveolar fibroblasts called alveolar fibroblast 1 (AF1) and alveolar fibroblast 2 (AF2) (also referred to as alveolar fibroblasts and pericytes, respectively)(*10*, *13*). AF1s expressed *Pdgfra*, *Limch1*, and *Tcf21* while AF2s express *Pdgfrb*, *Cox4i2*, and *Notch3* (Figure S1a,b). GO analysis revealed AF2s are enriched for vasculature niche support genes, suggesting these cells likely serve a “pericyte”-like function within the capillary plexus of the alveolus (Figure S1c). We used Pdgfra^CreERT2^ and Pdgfrb^CreERT2^ mouse lines to isolate AF1s and AF2s, respectively, and performed scRNAseq to characterize additional heterogeneity in these mesenchymal lineages (Fig. 1a). These data revealed that Pdgfra-traced cells labeled AF1s, adventitial, and peribronchial mesenchyme while Pdgfrb-traced cells labeled AF2s, adventitial, peribronchial, and vascular smooth muscle (VSMC) (Fig. 1b, c, Data S1, S2) We generated Pdgfra^H2B-GFP^:Pdgfrb^CreERT2^:Rosa26^LSL-tdTomato^ mice (Fig. 1d) and used immunohistochemistry (IHC) analysis to show that Pdgfra and Pdgfrb expression marked and segregated AF1s and AF2s, respectively, with a dual Pdgfra+/Pdgfrb+ labeled population only being observed around the airways and in the adventitial space (Fig. 1e, f, Data S3, S4).

**Figure 1:**
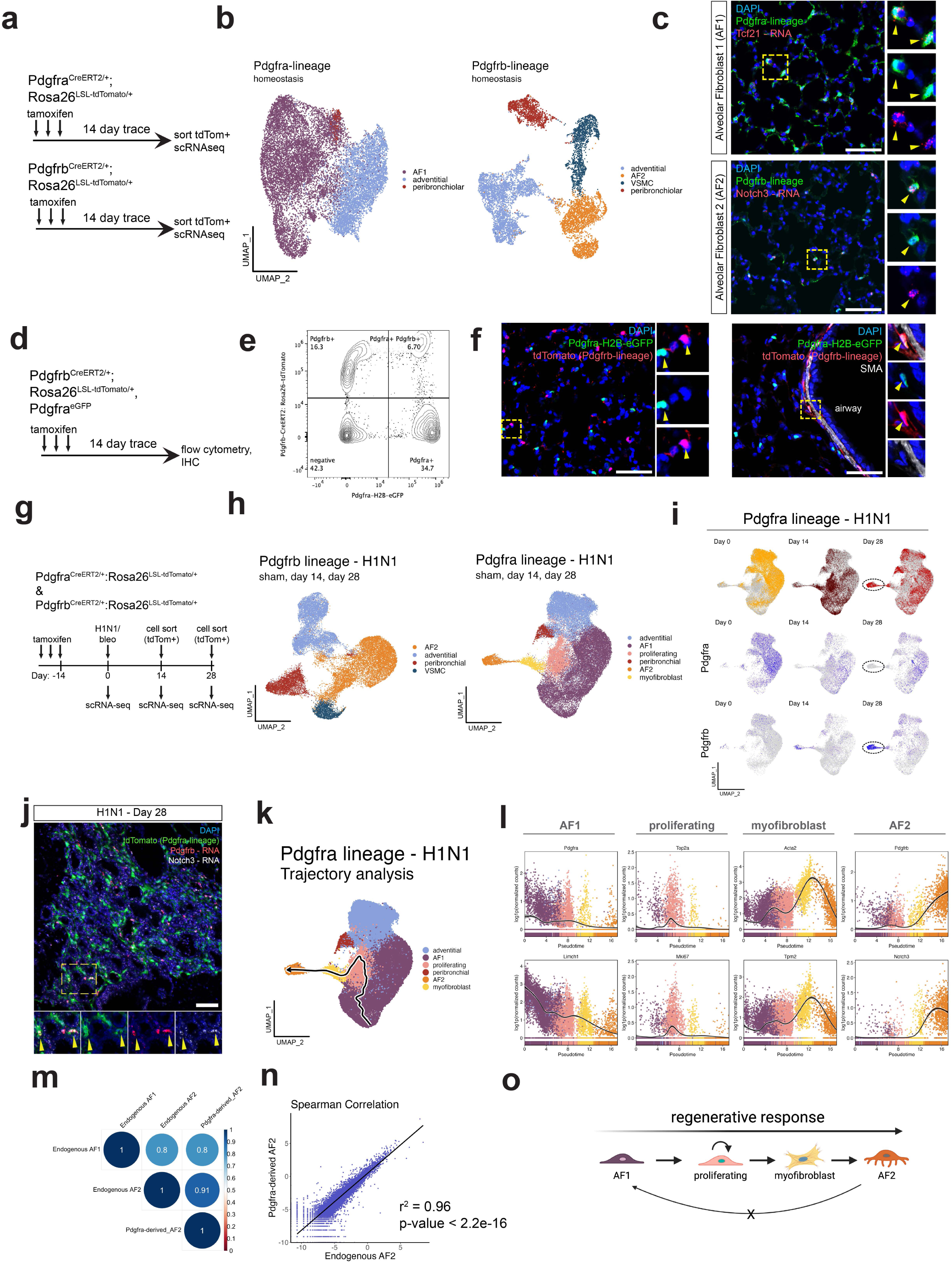
Reactivity and plasticity of the pulmonary Pdgfra+ mesenchymal cells in the lung. (a) experimental schematic of the approach to dissect Pdgfra+ and Pdgfrb+ cell heterogeneity in the adult mouse lung at homeostasis. (b) UMAP representation of the scRNA-seq data of the Pdgfra and Pdgfrb-lineage mesenchymal cells. (c) IHC and RNA in situ hybridization to visualize spatial location of AF1 and AF2 cells. (d) experimental schematic of the approach to dissect overlap of Pdgfra+ and Pdgfrb+ cells in the mouse adult lung. (e) flow-cytometry showing overlap and separation of Pdgfra+ (eGFP+) and Pdgfrb+ (tdTomato+) cells in the adult mouse lung. (f) IHC confirmation of flow-cytometry using Pdgfra and Pdgfrb as unique markers of AF1s and AF2s respectively. (g) schematic approach to study Pdgfra+ and Pdgfrb+ lineage mesenchymal cells after respiratory influenza and separately bleomycin-induced lung injuries. (h) UMAP representation of the scRNA-seq data of the Pdgfra and Pdgfrb-lineage mesenchymal cells after influenza infection. Data represents integrated libraries (sham, day 14, and day 28) for each lineage. (i) UMAPs and FeaturePlots showing division of Pdgfra and Pdgfrb-lineage mesenchymal cells at each timepoint, and the expression of *Pdgfra* and *Pdgfrb* of these cells over time. (j) IHC and RNA in situ hybridization for Pdgfra-derived AF2 cells (tdTomato+ Pdgfrb+ Notch3+) showing that these cells are embedded in iTCHs. (k) Slingshot trajectory analysis showing trajectory of AF1 cells differentiating into AF2 cells. (l) Dynamic gene expression through pseudotime of genes enriched in each cell state within the trajectory. (m) Correlation plot showing Spearman correlation coefficients of Pdgfra-derived AF2 cells compared to endogenous AF1 and AF2 cells. (n) Spearman correlation plot and calculation of R correlation coefficient and p-value revealing the high degree of similarity between wildtype AF2 and Pdgfra-derived AF2 cells. (o) summary schematic generated using Biorender.com

To determine the response of Pdgfra+ and Pdgfrb+ mesenchymal cells to lung injury, we performed lineage tracing with the Pdgfra^CreERT2^ and Pdgfrb^CreERT2^ mouse lines to fluorescently tag and trace Pdgfra/b+ cells and their future progeny and coupled this with a temporal series of scRNA-seq experiments across two pathologically relevant lung injury models: H1N1 influenza infection and bleomycin induced lung injury (Fig. 1g). Integration of Pdgfrb-lineage traced cells from all timepoints revealed these cells do not proliferate nor do these cells exhibit Pdgfrb-to-Pdgfra plasticity (Fig. 1h, S2a-d, S3a-d). In contrast, different subsets of Pdgfra-lineage cells emerged including a subset that exhibited a proliferative signature, another subset that exhibited a myofibroblast gene signature, and another subset that appeared to be AF2s differentiated from AF1s (Fig. 1h, S3e). Since Pdgfrb+ cells do not proliferate nor generate Pdgfra+ cells, these data indicate that the Pdgfra+ AF1 lineage, and not the adventitial or peribronchiolar lineages which express both Pdgfra and Pdgfrb, contribute to the Pdgfrb+ AF2 lineage. Indeed, transcriptional trajectory analysis suggested that the Pdgfra+ AF1s transit through a proliferative state before passing through the myofibroblast state and terminating in the AF2 state (Fig. 1k-l, S2h). These AF1-derived AF2s are apparent at day 28 after injury, express high levels of Pdgfrb, and lose expression of Pdgfra (Fig. 1i, S2f-g, S3f-g, Data S5). The AF1-derived AF2 cells are spatially localized within the severely damaged regions of the lung which contain dysplastic Krt5+ epithelium (Fig. 1j).

To determine in an unbiased manner whether Pdgfra-derived AF2s are transcriptionally similar to endogenous AF2s, we integrated the data from these cells with data from AF1s and AF2s isolated at homeostasis and performed Spearman correlation analysis. This analysis confirmed that Pdgfra-dervied AF2s exhibit a very high degree of similarity with endogenous AF2s (Fig. 1m, n). Together, these results indicate that after acute lung injury, AF1 cells act as mesenchyme progenitors, transiently proliferate, cross through a myofibroblast cell state, and then differentiate into Pdgfra-derived AF2s (Fig. 1o).

### Injury induced lung myofibroblasts are a transient state that arise from AF1 progenitors

Dynamic gene expression analysis across the trajectory of AF1s as they differentiate into AF2s suggests an increase in Pdgfrb+ expression that initiates during the transient myofibroblast state (Fig. 2a). To experimentally determine whether myofibroblasts are a transient state of AF1s while differentiating into AF2s, we examined whether myofibroblasts arise uniquely from the Pdgfra-lineage. Lineage tracing shows that AF1s, not AF2s, generated the vast majority of the Acta2+ (SMA+) cells after both H1N1 and bleomycin induced severe lung injury, consistent with previous reports (Fig 2b, c, S4a-c)(*14*). Lineage tracing was then performed at temporally defined times using both Pdgfrb^CreERT2^ and Acta2^CreERT2^ mice to tag and track mesenchymal cells as they transited through the various cell states before ending in AF2s. These studies revealed that lineage labeling Pdgfrb or Acta2 expressing cells during the time course of injury and regeneration captured the active myofibroblasts post-injury (Fig. 2d-h, S4d, e). By 90 days after injury, these previously labeled myofibroblasts extinguished Acta2 expression (Fig. 2i), retained Pdgfrb but not Pdgfra expression, and persisted in the severely damaged regions of the lung, directly adjacent to the Krt5-expressing dysplastic basal epithelium (Fig. 2j). These data demonstrate that the transient myofibroblast state induced by acute injury to the lung results in AF2 cells in persistently damaged regions of the lung.

**Figure 2:**
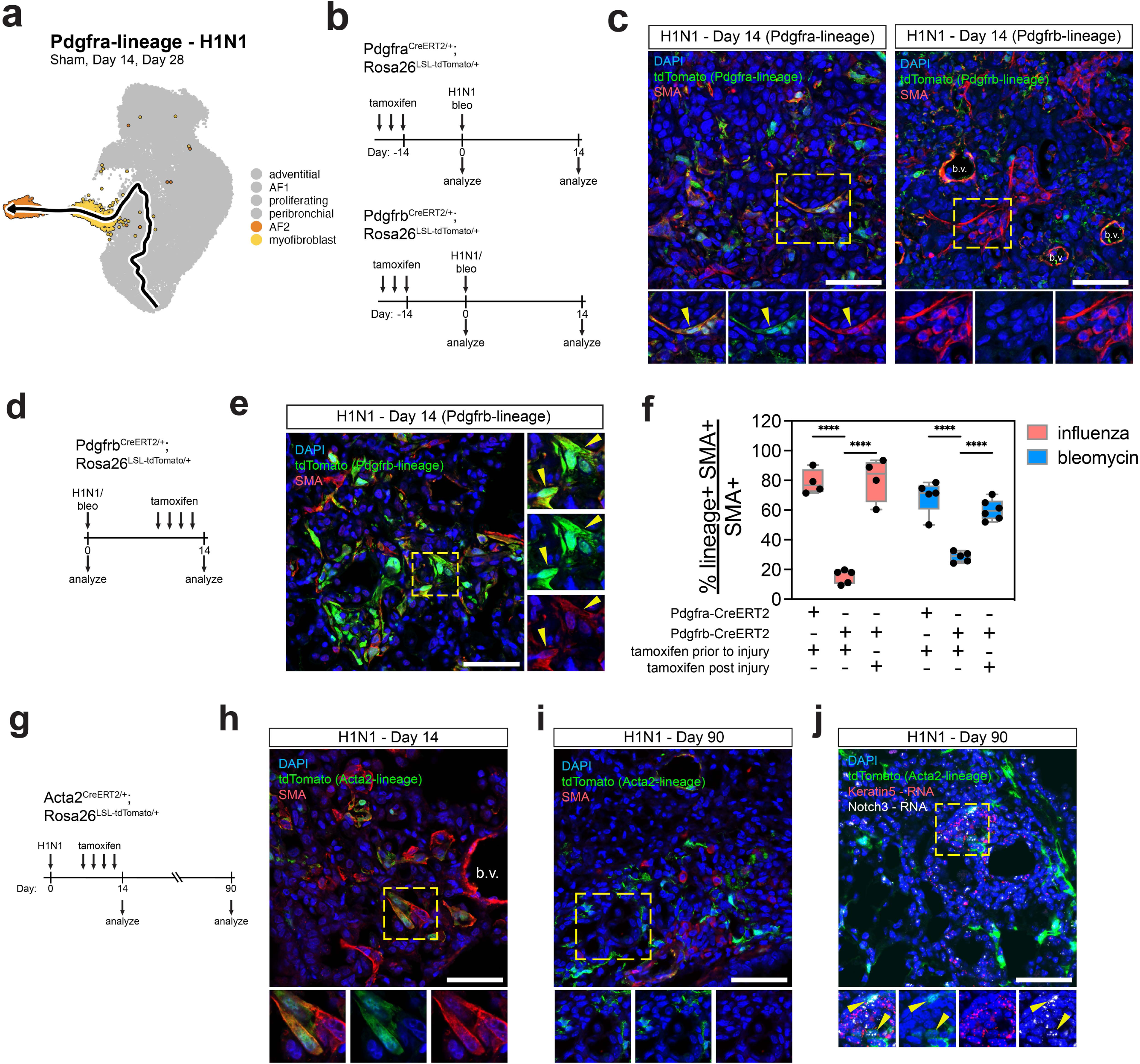
Source and fate of Acta2 expressing cells in the lung during injury and regeneration. (a) UMAP of scRNA-seq from Pdgfra-lineage cells after influenza (sham, day 14, and day 28) highlighting myofibroblasts and AF2 cells along the predicted trajectory of AF1 cells differentiating into AF2 cells. (b) Experimental schematic depicting approach to dissect whether Pdgfra+ or Pdgfrb+ cells are the primary producers of myofibroblasts in the lung. (c) IHC of the lineage reporter (tdTomato) and alpha smooth muscle actin (SMA) at 14 days after influenza infection showing that Pdgfra+ cells are the major producer of SMA+ myofibroblasts in the lung, relative to Pdgfrb+ cells. (d) Experimental schematic depicting approach to test whether lineage labeling Pdgfrb+ cells during injury labels SMA+ myofibroblasts. Mice were given daily tamoxifen from days 10-13 and collected for analysis at day 14. (e) IHC of the lineage reporter (tdTomato) and SMA at 14 days after influenza infection showing that labeling actively expressing Pdgfrb+ cells during injury captures the Pdgfra-derived SMA cells on their way to become AF2 cells. (f) Box plots showing quantification of the percentage of lineage-labeled SMA+ cells out of the total SMA+ cells after both influenza infection and separately bleomycin. (g) Experimental schematic depicting approach to capture and trace the Acta2 expressing cells during injury and after resolution. (h) IHC of the lineage reporter (tdTomato) and SMA showing that administration of tamoxifen after influenza infection labels the actively expressing Acta2+ myofibroblasts. (i) IHC showing previously labeled SMA cells extinguish SMA expression and reside in iTCHs. (j) IHC and RNA in situ hybridization showing that at later timepoints after influenza infection, labeled Acta2+ cells express the AF2 marker Notch3 and are embedded in iTCHs directly adjacent to the dysplastic Krt5-expressing epithelium. All scale bars represent 50 um. Each data point in (f) represents data obtained from an individual mouse (i.e. biological replicate). ****P<0.0001, evaluated by one-way ANOVA with Tukey’s adjustment for multiple comparisons.

### Blockade of AF1 cell division suppresses dysplastic repair and promotes euplastic epithelial regeneration

The localization of the Pdgfra-derived AF2s in the severely damaged regions of the lung containing dysplastic keratinized epithelium suggested that these cells may play a role in their emergence or persistence. We examined proliferation of AF1s given the scRNA-seq data suggesting these cells enter a transient proliferative phase during lung injury and observed a robust transient increase of proliferative Pdgfra-lineage labeled cells within the alveolar region starting at day 7 post-injury but little to no proliferation of the alveolar Pdgfrb-lineage cells (Fig. 3a-d). These data also showed an increase in the number of alveolar Pdgfra-lineage cells which was stable to at least 28 days post-injury (Fig. 3d). To determine the importance of the expansion of AF1 cells in alveolar regeneration, we genetically deleted Ect2, a Rho-GEF protein required for cytokinesis, in Pdgfra expressing cells to block cell division (herein termed Ect2^Pdgfra-KO^) (Fig. 3e, f). Cells lacking Ect2 can still undergo karyokinesis (nuclear division) but cannot undergo cytokinesis resulting in binucleation and a halt in cell division (Fig. 3f)(*15–17*). As expected, Ect2^Pdgfra-KO^ mutant cells became binucleated at 14 days post-injury, and these binucleated cells persisted until at least 28 days after injury (Fig. 3g). Surprisingly, blockade of Pdgfra+ cell division improved overall structure and reduced injury severity at 28 days after influenza infection, but not at 14 days after infection (Fig. 3h-j). These data suggest that Pdgfra+ cell division and expansion does not influence acute injury severity after influenza infection (i.e. 14 days after infection), but blocking this proliferative response allows the lung to engage euplastic regeneration after injury (i.e. 28 days after infection).

**Figure 3:**
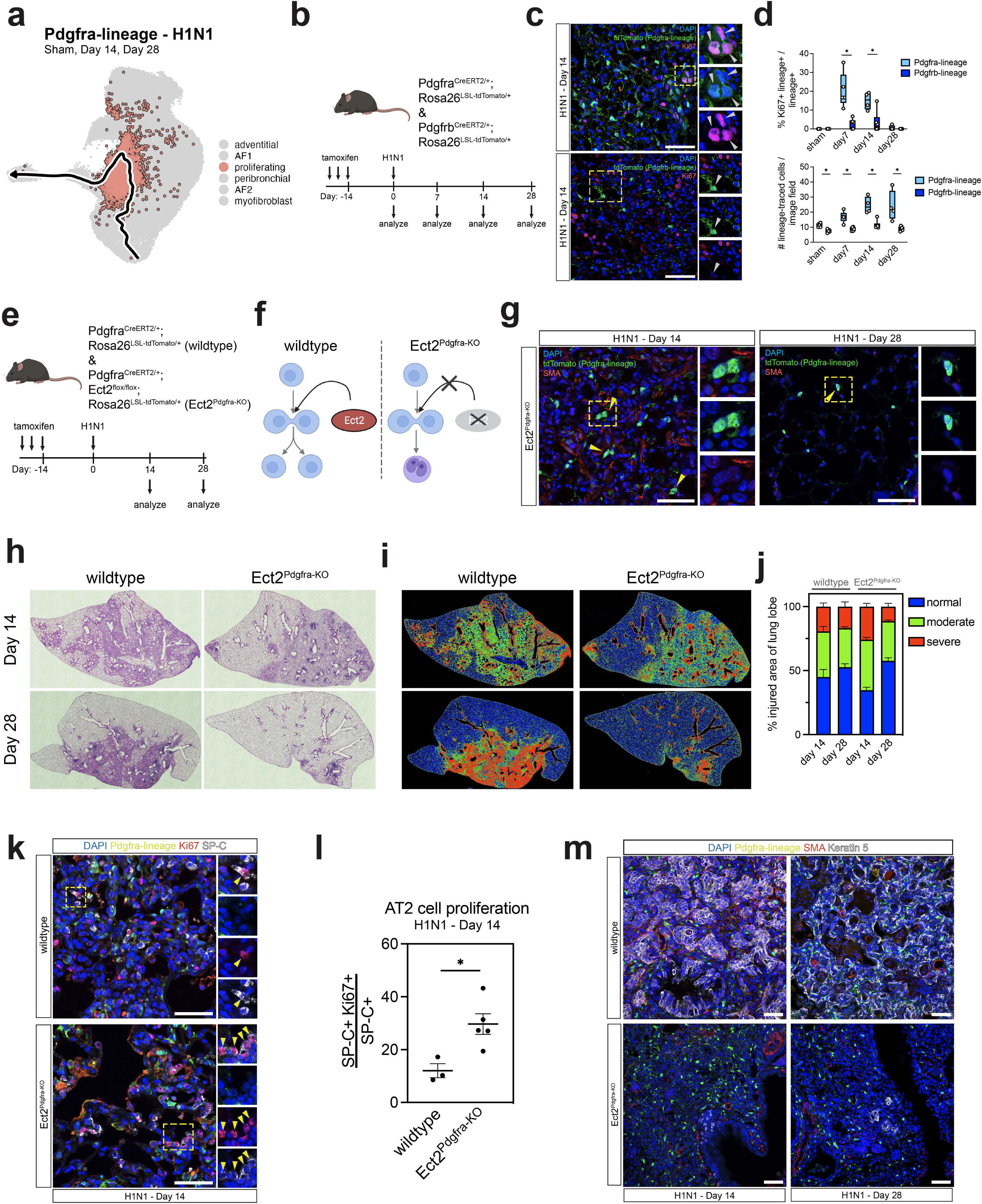
Selective inhibition of mesenchymal cell cytokinesis inhibits iTCH formation and promotes euplastic alveolar repair. (a) UMAP of scRNA-seq from Pdgfra-lineage cells after influenza (sham, day 14, and day 28) highlighting proliferating cells along the predicted trajectory of AF1 cells differentiating into AF2 cells. (b) Experimental schematic depicting approach to evaluate and quantifiy proliferation of Pdgfra+ and Pdgfrb+ mesenchymal cells over time after influenza infection. (c) IHC of the lineage-reporter (tdTomato) and Ki67 showing Pdgfra+ cell pool proliferates and expands after injury while Pdgfrb+ cells do not proliferate. (d) Box plots showing quantification of percentage of tdTomato+ Ki67+ cells out of the total tdTomato+ cells for both Pdgfra and Pdgfrb-lineage over time after influenza infection and also total number of lineage-traced cells per image field. (e) Experimental schematic depicting approach to evaluate the function of Pdgfra+ cell pool expansion after influenza injury via deletion of Ect2. (f) Cartoon schematic depicting function of Ect2 and the effect of deletion made with Biorender.com. (g) IHC showing bi-nucleated Pdgfra-lineaged trace cells after Ect2 deletion following influenza infection at days 14 and 28. (h) representative H&E sections of the left lobes after Ect2 deletion following influenza infection. (i) injury-severity algorithm showing regions of normal (blue), moderately damaged (green), and severely damaged (red) lung tissue after injury. (j) quantification of injury severities as a percentage of total lung area. (k) IHC of the Pdgfra-lineage reporter (tdTomato), SP-C, and Ki67 showing increased AT2 cells proliferation after Ect2 deletion in Pdgfra+ cells. (l) quantification of AT2 cell proliferation from IHC images. (m) IHC of Pdgfra-lineage (tdTomato), SMA, and Keratin5 showing absence of Krt5+ cells in the alveolar space after Ect2 deletion in Pdgfra+ cells. All scale bars represent 50 um. Each data point in panels d and l represents data obtained from a single mouse (i.e. biological replicate). n = 4-6 mice/group in panel j. All error bars represent SEM. *P<0.05, evaluated by un-paired t-test.

Since our histological analysis indicated that loss of AF1 cell expansion improved the overall lung architecture after injury, we assessed whether AT2 cell proliferation, a hallmark of alveolar regeneration, was affected in this model. Interestingly, blockade of AF1 cell expansion in Ect2^Pdgfra-KO^ mutants resulted in a significant increase in proliferation of AT2 cells at 14 days after injury compared to wildtype mice (Fig. 3k, l). Moreover, loss of AF1 cell expansion nearly eliminated the dysplastic Krt5+ cell response at 14 and 28 days after injury (Fig. 3m), demonstrating that expansion of AF1 cells is essential for dysplastic epithelial repair and suppresses the normal AT2 cell proliferative euplastic regeneration response after lung injury.

### AF1-derived AF2s form an <u>i</u>njury induced <u>t</u>issue ni<u>ch</u>e (iTCH) with dysplastic Krt5+ epithelium

To identify putative factors regulating the function of AF1-derived AF2s and how these cells may signal to Krt5+ dysplastic epithelium, transcription factor activity was assessed in these cells using DoRothEA, a computational tool which identifies putative transcription factors regulating cell population specific transcriptional networks (Fig. 4a-c, Data S6)(*18*). One of the top ranked transcription factors in the AF1-derived AF2s was Rbpj, the downstream transcription factor in the Notch signaling pathway (Fig. 4d). GSEA analysis demonstrated AF1-derived AF2s are strongly enriched in expression of Notch signaling pathway components (Fig. 4e), including Notch receptors (e.g. *Notch1, 2, 3*) and downstream target genes (*Heyl, Hey2, Ccnd1, Postn, Tbx2*) (Fig. 4f). Spatial analysis of the Notch “active” Pdgfra cells assessed by intracellular Notch1-ICD immunostaining(*19*), revealed these cells are located in the dysplastic alveolar niche after injury, directly adjacent to the Krt5+ epithelium (Fig. 4g).

**Figure 4:**
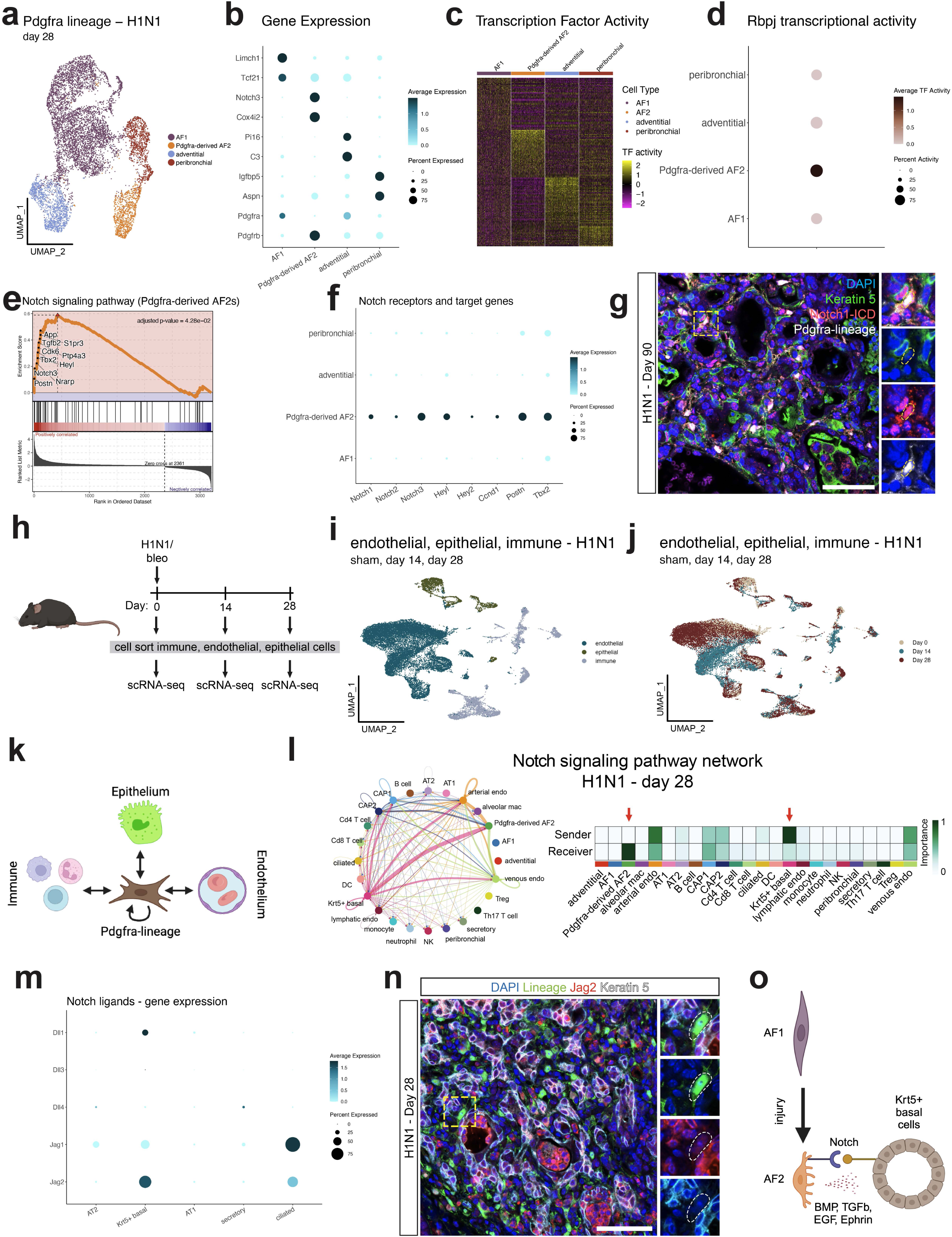
Pdgfra-derived AF2 cells are embedded within iTCHs and are enriched for Notch signaling. (a) UMAP of scRNA-seq data from Pdgfra-lineage cells at 28 days after influenza infection. (b) DotPlot showing unique gene experession markers of each cluster shown in panel (a). (c) Heatmap depicting the top 25 active transcription factors unique to each cell cluster. (d) DotPlot showing activity of Rbpj, the main transcription factor downstream of Notch signaling, in each cell cluster shown in panel (a) demonstrating that Pdgfra-derived AF2 cells have the highest Rbpj activity relative to other cell types. (e) GSEA analysis showing Pdgfra-derived AF2 cells are enriched for downstream target genes of Notch signaling. (f) DotPlot showing expression level of Notch receptors and downstream target genes in cell clusters shown in panel (a). (g) IHC at 90 days after influenza infection showing Notch+ Pdgfra-lineage cells are embeded within iTCHs directly adjacent to the dysplastic Krt5+ epithelium. (h) Experimental schematic depicting approach to temporally profile immune, endothelial, and epithelial cells using scRNA-seq after influenza infection and separately bleomycin. (i) UMAP representation of the scRNA-seq data showing endothelial, epithelial, and immune cells. (j) UMAP representation of the scRNA-seq data showing aggregation of cells by time point (sham, day 14, and day 28). (k) Cartoon schematic showing approach to examine which cell types are communicating with Pdgfra-lineage cells after injury made with Biorender.com. (l) Notch signaling network at 28 days after influenza infection showing a unique niche between the Krt5+ dysplastic basal cells, the primary sender of the Notch signals, and Pdgfra-derived AF2 cells, the primary receiver of Notch signals. (m) DotPlot showing expression of Notch ligands across the distinct epithelial cell types at day 28 after influenza infection showing Krt5+ basal cells express high levels of Dll1 and Jag2. (n) IHC showing Jag2+ Krt5+ dysplastic epithelial cells exist directly adjacent to the Pdgfra-lineage mesenchymal cells at day 28 after influenza infection. (o) Cartoon summary schematic made with Biorender.com. All scale bars represent 50 um.

To identify which cell is producing the Notch ligands, we performed scRNA-seq on epithelial, endothelial, and immune cells after both H1N1 influenza infection and bleomycin induced injury at days 14 and 28 (Fig. 4h-j). We then performed CellChat analysis to identify proper sending-receiving cell patterns induced upon acute lung injury (Fig. 4k)(*20*). These data showed that Krt5+ dysplastic epithelial cells are the primary producer of Notch ligands, with the AF1-derived AF2s as the primary Notch receiver (Fig. 4l, m, S5a-c). Further analyses revealed that AF1-derived AF2 cells send BMP, TGFb, EGF, and Eph-Ephrin signaling back to the Krt5+ epithelium (Fig. S5b). IHC additionally revealed that Jag2-expressing Krt5+ dysplastic basal cells are directly adjacent to the AF1-derived AF2s 28 days after injury (Fig. 4n). These analyses demonstrate that AF1-derived AF2s exhibit active Notch signaling and generate a unique injury induced tissue niche or iTCH with the Krt5+ dysplastic epithelium (Fig. 4o).

### Lineage selective inhibition of mesenchymal Notch signaling blocks dysplastic repair and promotes euplastic alveolar regeneration

To determine whether Notch signaling in AF1-derived AF2s is critical for formation of iTCHs, Pdgfra^CreERT2^:Rosa26^dnMAML^ mice (Notch^Pdgfra-KD^) were used to block intracellular Notch signaling in AF1 cells. Notch^Pdgfra-KD^ mice were subjected to influenza injury and examined at 14 days post-injury (Fig. 5a). Blocking intracellular Notch signaling within AF1s improved overall survival after influenza infection (Fig. 5b). Quantitative histological analyses from lungs of Notch^Pdgfra-KD^ mice revealed a dramatic reduction in iTCHs relative to wildtype mice post-influenza injury (Fig. 5c, d). Remarkably, this reduction in dysplastic repair correlated with an increase in the number of AT2 cells throughout the remaining damaged regions of the lung, indicating that suppression of mesenchymal intracellular Notch signaling rewires cell-cell communication in the alveolus to favor euplastic regeneration and reduce iTCH formation (Fig. 5e). CD45+ immune cell infiltrate was still observed in the severely damaged regions of the lung at 14 days post injury (Fig. 5f). To further examine whether mesenchymal intracellular Notch signaling regulates persistence of iTCHs after influenza, micro-computed tomography (uCT) was performed on mice 90 days after influenza infection. Locations of iTCHs were based on aerated lung volume analysis. Control influenza infected wildtype mice exhibited a significant reduction in aerated lung volume at 90 days compared to uninfected mice (Fig 5g, h), consistent with prior work demonstrating a reduction in pulmonary gas-exchange function in mice one year after a single influenza infection(*8*). Blocking Notch signaling in the AF1 lineage inhibited the reduction of aerated lung volume, further indicating that Notch activity in these mesenchymal cells is essential for promoting dysplastic repair over euplastic regeneration in the lung (Fig. 5g, h).

**Figure 5:**
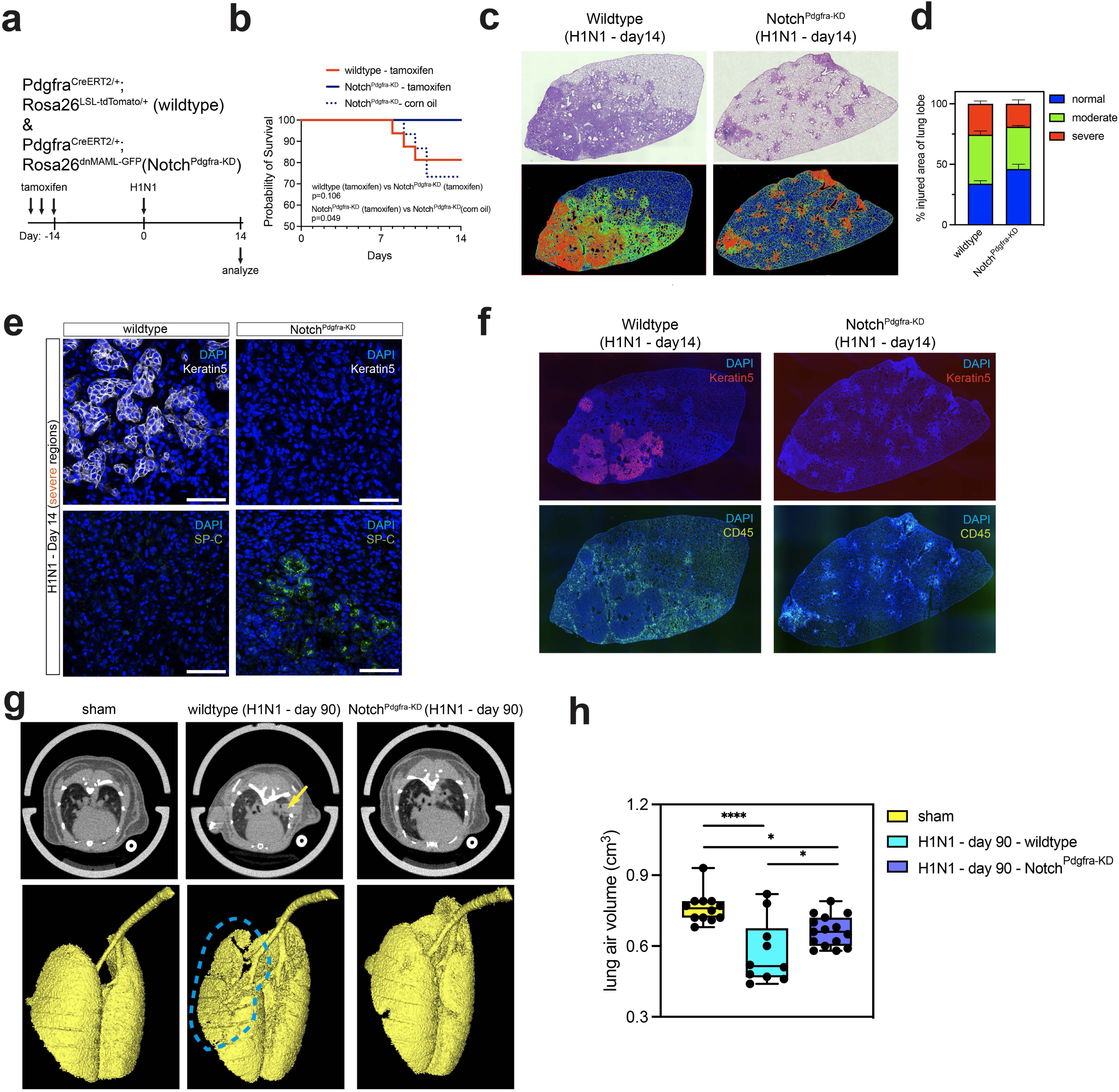
Inhibition of mesenchymal Notch signaling blunts dysplastic alveolar remodeling. (a) Experimental schematic showing approach to understand the function of intracellular Notch signaling in Pdgfra+ cells. (b) Survival curve of wildtype and Notch^Pdgfra-KD^ mice after influenza infection. P-values calculated from Log-rank (Mantel-Cox) tests. (c) H&E and injury severity zones of the left lobes from wildtype and Notch^Pdgfra-KD^ mice after influenza infection. (d) quantification of injury severity zones from wildtype and Notch^Pdgfra-KD^ mice. (e) IHC showing that wildtype mice iTCHs contain Krt5+ dysplastic epithelium and no SP-C+ AT2 cells, but in Notch^Pdgfra-KD^ mice iTCHs do not contain Krt5+ epithelium with an increase in SP-C+ AT2 cells. (f) whole lobe IHC analysis showing remain iTCHs in Notch^Pdgfra-KD^ mice exhibit dense CD45+ immune cell infiltration. (g) uCT scans of sham, wildtype (day 90 after influenza infection), and Notch^Pdgfra-KD^ (day 90 after influenza infection) showing a reduction in gross lung scar tissue formation in Notch^Pdgfra-KD^ compared to wildtype mice at 90 days after influenza infection. (h) Box plot of uCT analysis showing a rescue in total lung air volume in Notch^Pdgfra-KD^ (day 90 after influenza infection) relative to wildtype (day 90 after influenza infection). All scale bars represent 50 um. All error bars represent SEM. Each data point in (h) represents data obtained from an individual mouse (i.e. biological replicate). n = 5-6 mice/group in panel d. ****P<0.0001, *P<0.05, evaluated by one-way ANOVA with Tukey’s correction for multiple comparisons.

Since Notch signaling is also observed in the dysplastic Krt5+ epithelium within iTCHs (Fig. 4g), Notch signaling was inactivated within this basal cell derived epithelium during influenza induced lung regeneration using Krt5^CreERT2^:Rosa26^dnMAML^ mice (here after called Notch^Krt5-KD^) (Fig. S6a). In contrast to inactivating intracellular Notch within the Pdgfra+ cells, inactivating intracellular Notch in the Krt5+ dysplastic epithelium (Notch^Krt5-KD^) did not improve survival after influenza injury (Fig. S6b). IHC analysis showed Krt5+ dysplastic epithelium continued to egress into the severely damaged alveolar region upon cell autonomous loss of Notch signaling (Fig. S6c). We next employed *in vitro* assays to assess Notch^Krt5-KD^ basal cell behavior. We first confirmed that expression of dnMAML in these cells *in vitro* strongly inhibited Notch signaling (Fig. S6d). However, genetic inactivation of Notch signaling did not have any effect on cell proliferation or organoid growth, in stark contrast to chemical gamma-secretase inhibition with DBZ, which was previously shown to inhibit basal cell growth(*7*) (Fig. S6e-i). Collectively, these data demonstrates that intracellular mesenchymal Notch signaling is the dominant mechanism which controls the generation of the dysplastic repair response and iTCH formation after acute lung injury.

### Blockade of iTCH formation via Notch inhibition increases flow and direction of Wnt and Fgf signaling in the alveolar niche

To identify the mechanism by which blocking Notch signaling within mesenchymal cells rewires the alveolar niche to support euplastic over dysplastic responses, we performed scRNA-seq on whole lungs from wildtype and Notch^Pdgfra-KD^ mice at 14 days post-influenza infection (Fig. 6a-c, S7a-e). These data revealed three distinct AF1 cell populations which we named AF1a, b, and c (Fig. 6d). While AF1a was composed of relatively equal percentages of cells derived from wildtype and Notch^Pdgfra-KD^ lungs, AF1b was strongly enriched for AF1 cells derived from wildtype mice, while AF1c was strongly enriched for AF1s derived from Notch^Pdgfra-KD^ mice (Fig. 6e). None of the remaining mesenchymal cell clusters were dominated by either wildtype or Notch^Pdgfra-KD^ cells (Fig. 6e), suggesting that inactivating intracellular Notch within the Pdgfra-lineage preferentially regulates the AF1 cell response after influenza injury. We next performed pathway analysis using the genes uniquely enriched in either the AF1b or AF1c cell clusters. We found that AF1b was enriched in gene expression for ECM-receptor interactions, proteoglycans, and focal adhesions (Fig. 6f). The AF1c cluster dominated by Notch^Pdgfra-KD^ cells was enriched in gene expression of MAPK, TNF, and PI3K-Akt signaling pathways and genes such as *Fgf7*, *Vegfa*, and *Wnt2* (Fig. 6f). These data indicate that inactivation of Notch signaling in the Pdgfra-lineage blunts the fibrotic and myofibroblast response and upregulates euplastic regenerative signaling patterns in support of AT2 cell proliferation and survival including Wnt and Fgf(*11*, *16*, *21–23*). Consistent with our IHC analyses in Figure 5e-f, the scRNA-seq analysis of the lung epithelium in Notch^Pdgfra-KD^ animals revealed a stark reduction of Krt5+ basal cells in the Notch^Pdgfra-KD^ dataset, with an increase in total AT1 and AT2 cells (Fig. 6g, h).

**Figure 6:**
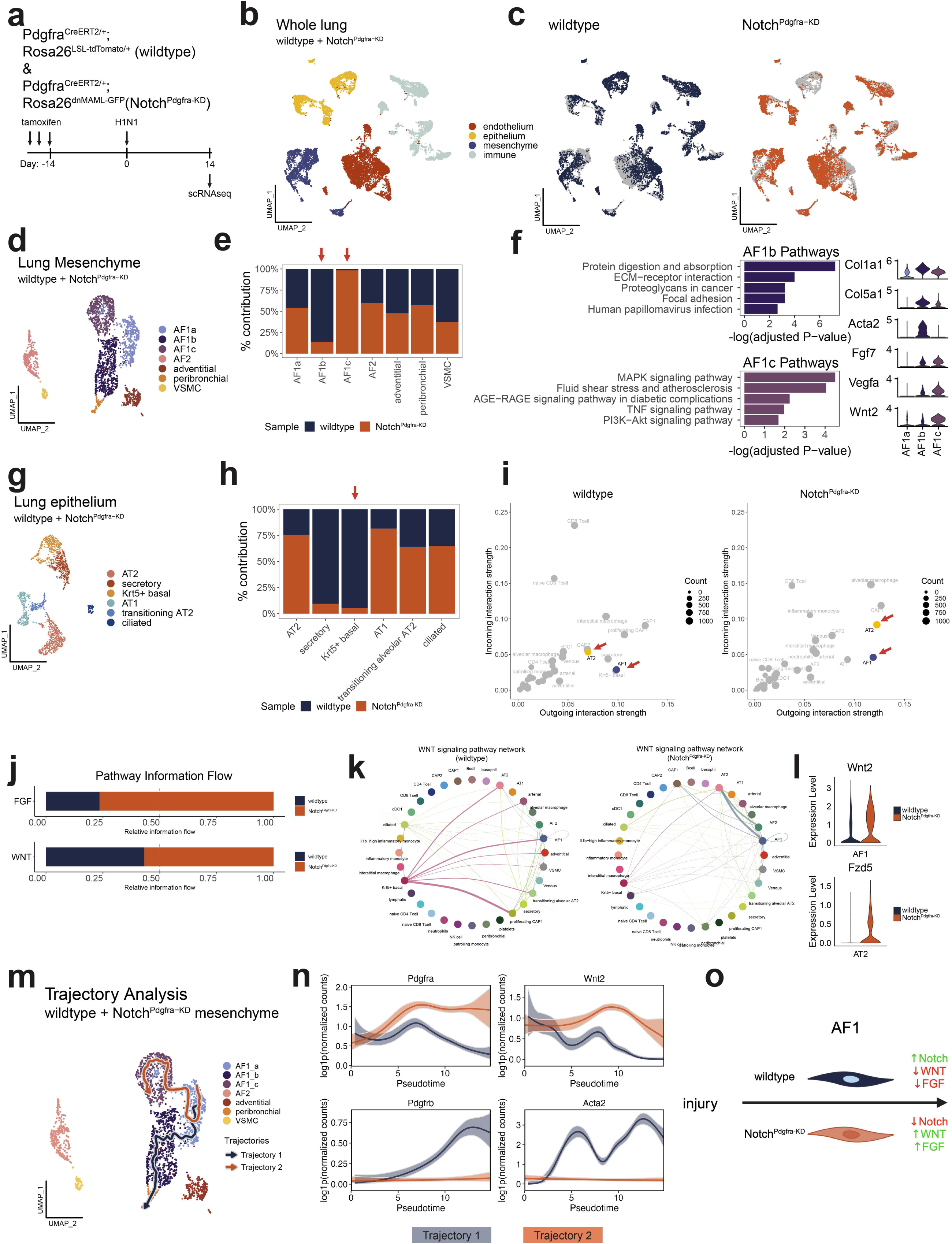
Mesenchymal Notch signaling rewires the alveolar signaling niche. (a) Experimental schematic showing approach to identify the mechanism by which inhibiting intracellular mesenchymal Notch signaling rescues euplastic alveolar regeneration. (b) UMAP of scRNA-seq data from wildtype and Notch^Pdgfra-KD^ Pdgfra-lineage cells at 14 days after influenza infection. (c) UMAP highlighting spatial location of cells from each condition (wildtype and Notch^Pdgfra-KD^). (d) UMAP of mesenchymal cells showing three distinct clusters of AF1 mesenchymal cells. (e) Contribution of each condition (wildtype and Notch^Pdgfra-KD^) to the cell clusters shown in panel (d) showing AF1b is enriched for wildtype AF1 cells and AF1c is enriched for Notch^Pdgfra-KD^ AF1 cells. (f) KEGG pathway analysis of the gene expression programs enriched in AF1b and AF1c clusters and violin plots of relevant genes from the highlighted pathways. (g) UMAP of epithelial cells. (h) Contribution of each condition (wildtype and Notch^Pdgfra-KD^) to each cell cluster shown in panel (g) showing strong reduction in Krt5+ basal cells after Notch inactivation in the Pdgfra+ cells. (i) Incoming/outgoing signaling strength of all identified celltypes from panel (b) in both conditions (wildtype and Notch^Pdgfra-KD^) showing an increase in the incoming signaling from AT2 cells and an increase in outgoing signaling in AF1 cells in Notch^Pdgfra-KD^ mice compared to wildtype mice. (j) Differential pathway information flow for FGF and WNT signaling showing an overall system wide increase in FGF and WNT signaling in Notch^Pdgfra-KD^ mice compared to wildtype mice. (k) WNT signaling network in wildtype and Notch^Pdgfra-KD^ conditions showing a re-wiring of the WNT signaling niche after Notch inactivation in the Pdgfra+ cells. (l) Violin plots showing an upregulation of Wnt ligands (Wnt2) and receptors (Fzd5) in AF1 and AT2 cells respectively in Notch^Pdgfra-KD^ mice compared to wildtype. (m) Slingshot trajectory analysis depicting potential trajectory of AF1c cells, the common AF1 subcluster between wildtype and Notch^Pdgfra-KD^ mice, transiting into AF1b (dominated by wildtype) or AF1c (dominated by Notch^Pdgfra-KD^) subclusters. (n) Dynamic gene expression from each trajectory shown in panel (m) of AF1 markers (*Pdgfra*, *Wnt2*) and transient AF1 markers on their way to becoming AF2 cells (*Pdgfrb*, *Acta2*). (o) Cartoon summary schematic generated with Biorender.

We next examined differential signaling networks between wildtype and Notch^Pdgfra-KD^ cells that could explain the rewiring of the iTCHs upon loss of mesenchymal Notch signaling. This analysis showed an overall increase in outgoing signals emanating from AF1s and an increase in both outgoing and incoming signals into AT2s in Notch^Pdgfra-KD^ mutants (Fig. 6i). Cell type specific analysis revealed an increase in AF1-to-AT2 communication after Notch inactivation (Fig. S8a, b). Pathway specific analysis revealed an increase in Fgf and Wnt signaling in Notch^Pdgfra-KD^ mutants compared to wildtype (Fig. 6j, S8c). In control mice, the dysplastic Krt5+ epithelium was the dominant sender of Wnt signaling. However, in Notch^Pdgfra-KD^ mutants the Wnt signaling network is rewired with the new dominant Wnt signaling relationship being between AF1 and AT2 cells (Fig. 6k). Consistent with this new signaling relationship, we observed an increase in Wnt ligand expression in AF1s including *Wnt2*, and an increase in Wnt receptor expression in AT2s including the pro-regenerative receptor *Fzd5* in Notch^Pdgfra-KD^ mutants (Fig. 6l)(*24*). Furthermore, we observed a reduction in BMP, FGF, and Gas signaling from AF1 to Krt5+ dysplastic basal cells after injury demonstrating that intracellular Notch signaling in the mesenchyme directs the signaling network within the alveolus and can be targeted to promote epithelial euplastic repair (Fig. S8d). These data reveal a change in Wnt and Fgf signaling flow and direction in the alveolus depending on the Notch signaling status in AF1 cells.

To determine whether blocking intracellular Notch signaling in the Pdgfra-lineage prevents AF1-to-AF2 differentiation, we performed Slingshot trajectory analysis on the lung mesenchyme in control and Notch^Pdgfra-KD^ mutant mice resulting in two primary trajectories (Fig. 6m). Trajectory 1 traveled from AF1a through AF1b, the AF1 subcluster dominated by wildtype cells (Fig. 6e) and exhibited a loss of AF1 markers (*Pdgfra* and *Wnt2*), with a concordant increase of AF2 and myofibroblast markers (*Pdgfrb* and *Acta2*) suggesting these cells are transiting to becoming AF2s (Fig. 6n). In contrast, trajectory 2 traveled from AF1a through AF1c, the AF1 subcluster dominated by Notch^Pdgfra-KD^ mutant cells (Fig. 6e). Dynamic gene expression through trajectory 2 revealed that these cells maintained high expression of AF1 markers (*Pdgfra* and *Wnt2*) demonstrating that after influenza injury, AF1 cells transit to become AF2 cells and that this decision is dependent on Notch resulting in opposing signaling by Wnt and Fgf pathways (Fig. 6n). Together, these data show that loss of Notch signaling in AF1s leads to their reprogramming and shunting into a pro-euplastic regenerative phenotype, which inhibits dysplastic lung repair (Fig. 6o).

### Segregation of human lung disease phenotypes by the presence of iTCHs and rewiring of alveolar signaling flow

Diseases including post-COVID-19 lung disease is characterized by extensive myofibroblast cell expansion and fibrosis(*25*). Moreover, bleomycin-induced lung injury in humans is a rare form of reactive lung injury that additionally results in fibrosis(*26*). Phenotypes in both of these diseases are initiated by acute lung injury. In contrast, chronic lung diseases such as chronic obstructive pulmonary disease (COPD) are characterized by a degenerative emphysematous phenotype leading to a loss of alveolar structure, which is also exhibited by Alpha 1 anti-Trypsin deficiency (AAT), a genetic form of COPD(*27*). The role of dysplastic epithelial responses in these diseases is not well established, although the presence of KRT5+ epithelium in the alveolar compartment has been reported in a variety of fibrotic lung diseases(*8*).

To determine whether the rewiring of the alveolar signaling crosstalk by iTCHs could be used to segregate different phenotypes of human lung disease, a scRNA-seq dataset from bleomycin induced human lung disease and Alpha-1 anti-Trypsin human lung disease were newly generated and combined with previously published post-COVID-19 fibrotic lung disease datasets, to determine whether pathologic alveolar signaling networks driven by iTCHs are present in these diseases. Consistent with the mouse mesenchyme data, analysis of human mesenchymal cells from healthy control lungs revealed 5 distinct mesenchymal cell populations including the two distinct populations of alveolar mesenchymal fibroblasts seen in mice: AF1s (*PDGFRA*+ *TCF21*+) and AF2s (*PDGFRB*+ *NOTCH3*+) (Fig. S9a-d, Data S7). To determine whether AF2s and dysplastic basal cells communicate to form an iTCH in human post-COVID-19 and bleomycin induced human lung disease, single cell data was analyzed to examine ligand-receptor interactions between the epithelial and mesenchymal cell populations. These data revealed a robust NOTCH niche in the post-COVID-19 and bleomycin human lungs between the AF2s and the KRT5+ basal cells (Fig. 7a). This is supported by IHC showing AF2s, but not AF1s, are present adjacent to KRT5+ dysplastic epithelial cells in post-COVID-19 fibrosis and acute bleomycin injury (Fig. 7b).

**Figure 7:**
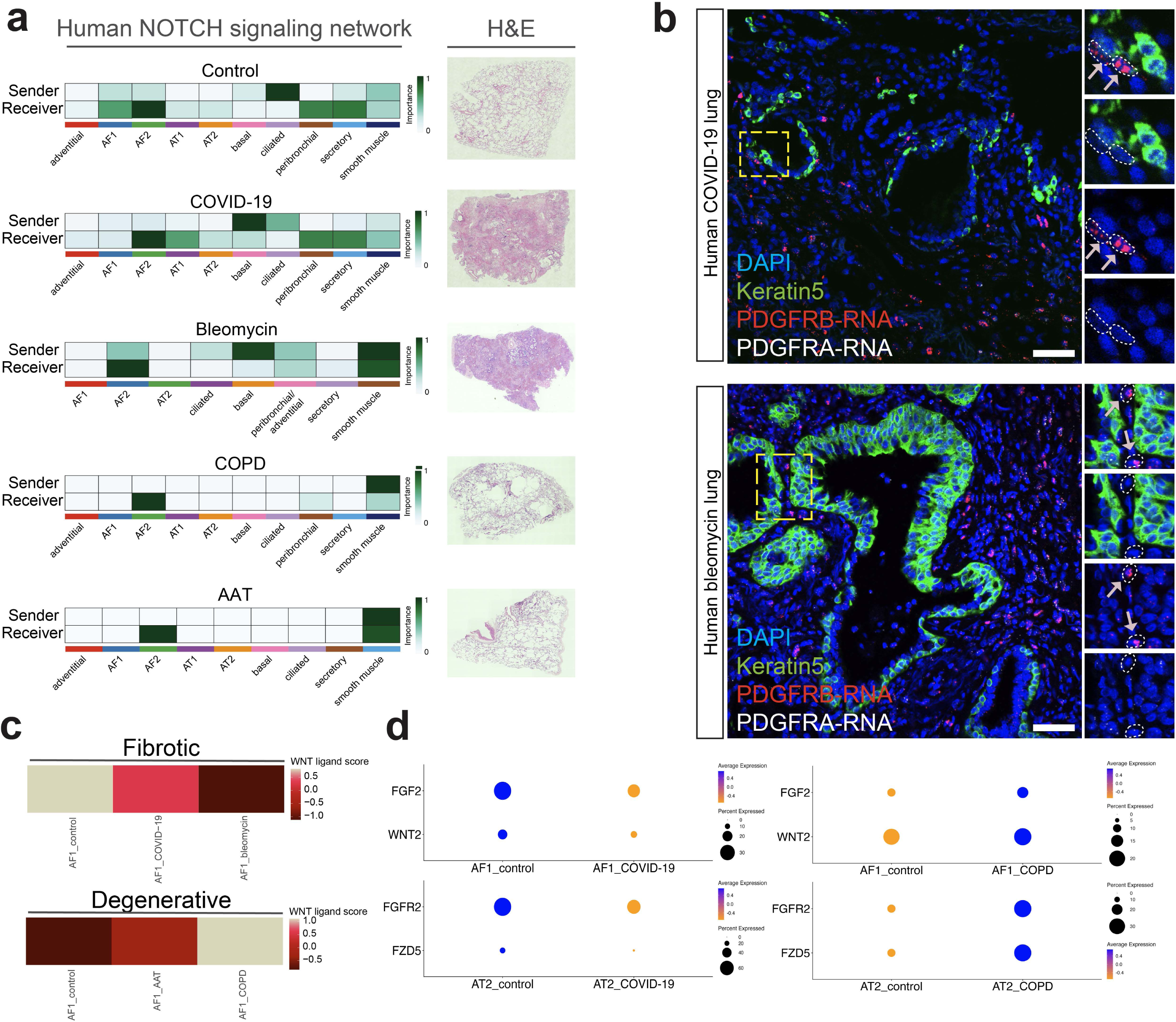
Phenocopying chronic human lung diseases using NOTCH, WNT, and FGF mesenchymal-epithelial signaling networks. (a) Notch signaling network between mesenchymal and epithelial cells from a scRNA-seq mutli-disease dataset of a variety of human lung diseases along with representative H&E tile scans of the distal lung isolated from each disease. (b) IHC and RNA in situ hybridization showing PDGFRB+ mesenchymal cells exist directly adjacent to KRT5+ dysplastic epithelium in COVID-19 and bleomycin-induced human lung injuries. (c) WNT ligand module score in AF1 cells from fibrotic (COVID-19 & bleomycin) human lungs and degenerative (AAT & COPD). (d) DotPlots showing relative expression level of FGF and WNT ligands and receptors within AF1 and AT2 cells from control, COVID-19, and COPD human lungs.

In contrast to the fibrotic lung diseases, we observed an increase in WNT ligand expression in AF1 cells from both COPD and AAT lungs compared to normal donor lungs, suggesting a reversal of a NOTCH/WNT axis in degenerative versus fibrotic alveolar diseases (Fig. 7c). This is reflected by a reduction in overall WNT signaling with decreased expression of FGF and WNT ligands (i.e. FGF2 and WNT2) in AF1s and receptors (i.e. FGFR2 and FZD5) in AT2s in fibrotic diseases (Fig. 7d). This is in stark contrast to COPD and AAT lung disease, where a rewiring of the NOTCH mediated signaling niche occurs and KRT5+ cells are no longer the dominant sender of the NOTCH ligands (Fig. 7a). Moreover, COPD and AAT do not exhibit dysplastic iTCH formation (Fig. 7a). Remarkably in COPD and AAT, there is an increase in FGF and WNT ligands and receptors in AF1 and AT2 cells, mimicking the euplastic regenerative response in the mouse lung upon loss of Notch signaling in AF1 derived AF2s and suggesting that in these degenerative diseases, a euplastic regeneration response is being engaged albeit inefficiently (Fig. 7d). These findings demonstrate a unique injury-induced tissue niche is formed between the mesenchyme and dysplastic epithelium in human fibrotic but not degenerative lung diseases, revealing that the presence and action of iTCHs balances the dysplastic versus euplastic response is species conserved in the mouse and human lung.

## Discussion

The cellular and architectural complexity of the respiratory system has made it difficult to unravel the specific cell lineages responsible for promoting functional lung regeneration. Our study reveals that the bifurcation decision between euplastic regeneration and dysplastic repair is governed by Notch signaling in a specific mesenchymal lineage called AF1 cells, which form an injury-induced niche or iTCH, resulting in a maladaptive structure which does not effectively participate in gas exchange. The AF1 response to acute injury involves transiting across multiple cell states including proliferation and myofibroblast, before assuming an AF2 phenotype, which is characterized by high Notch signaling activity. This AF1-AF2 plasticity and expansion is highjacked by Krt5+ dysplastic epithelial cells invading the alveolar niche to establish the iTCH. iTCH signaling patterns can be used to phenotype human lung diseases into either acute fibrotic responses such as COVID-19 or bleomycin induced acute injury, or degenerative phenotypes such as COPD and AAT deficiency. This phenotypic switch is characterized by altered signaling flows of hallmark regenerative pathways including WNT and FGF in the alveolar niche. This study reveals an emergent signaling niche in injured and diseased lungs that defines the tissue logic deciding between euplastic regeneration or dysplastic repair providing critical insight into phenotyping human lung disease responses.

The role of mesenchymal-epithelial signaling in organ development has been well described(*28–30*). In the lung, changes in phenotypes within the mesenchymal lineage occurs throughout development with the early lung mesenchyme being derived from a Wnt2+/Pdgfra+ progenitor cell(*28*). Our study demonstrates that Pdgfra+ alveolar mesenchymal cells in the adult lung called AF1 cells retain their developmental capacity to regenerate and replenish the non-proliferative Pdgfrb+ AF2 cell pool after injury. The current study shows that the dysplastic Krt5+ epithelium hijacks this AF1 cell plasticity, using a paracrine signaling network instigated by intracellular mesenchymal Notch signaling to form an iTCH. While some previous work has suggested Notch signaling within the basal cell lineage as a driving factor in formation of Krt5+ dysplastic epithelium(*7*), our current study demonstrates that Notch signaling in AF1 cells is the dominant driver of this phenomena to control iTCH formation. A previous study also suggests that Gli1+ mesenchyme can regulate the dysplastic epithelial response due to acute injury(*31*).

The Krt5+ cells that invade the alveolar space of the lung after acute injury are derived from rare intrapulmonary basal stem cells in the mouse(*7*, *9*). Their ability to rapidly expand and migrate into a completely different compartment of the lung and establish an iTCH in partnership with AF2s derived from AF1s, demonstrates the potent ability of basal stem cells to respond to an acute insult and the necessity of these cells to organize as a Notch dependent niche. This iTCH invasion competes with the euplastic regenerative response which is governed by AT2 cell proliferation and AT2-AT1 differentiation after acute injury(*4*, *5*). The balance between the airway and alveolar reactions to injury and certain lung diseases is an example of a “stem cell collision” that leads to the rewiring of the alveolar niche and a long-term reduction of gas exchange function. Balancing these responses is central to developing new therapies to combat long term consequences of acute respiratory injury such as after COVID-19 and influenza infections.

The ability of AF1 cells to differentiate into AF2 cells likely plays a critical role in replacing lost AF2s upon injury or during homeostatic turn-over since AF2s do not proliferate in response to injury. Severe injury can cause an imbalance in multiple processes that may not occur in more moderate injuries, including significant loss of AT1 and AT2 cells due to viral-induced cytotoxic cell death in influenza infection or bleomycin-induced acute pulmonary toxicity. This may lead to a larger degree of AF1 proliferation and AF1-AF2 differentiation than in less severe injuries which do not result in a dramatic loss of alveolar epithelium. Denuding the alveolar epithelium would provide the invading Krt5+ basal cells an open environment to establish themselves, driving iTCH formation. The high level of iTCHs in both COVID-19 and bleomycin acute injured lungs indicates that this process is conserved in acute injury in the human lung.

Repetitive acute injuries may underlie the emergence or progression of chronic lung disease(*32*, *33*). Most of the respiratory system is quiescent at homeostasis but reacts rapidly to injury, leading to the concept that it is a facultatively regenerative organ. While AT2 cells are well characterized as the resident alveolar epithelial stem/progenitor cell, they have critical roles in normal homeostatic lung function including surfactant production and recycling as well as an important immune sensing role(*1*, *34*). AT2 progenitor activity is active at only a very low levels during homeostasis where they slowly self-renew and differentiate into AT1 cells(*16*). In contrast, basal cells are a dedicated stem cell lineage whose primary purpose is to harbor the cellular and genetic information for regenerating and replacing all airway epithelial lineages during homeostatic turn over and after injury(*4*). Given the dedicated nature of basal stem cells, it is not surprising that when activated the basal cells out compete AT2 cells for repopulating severely damaged regions of the lung. Targeted approaches which suppress overactive basal cell behavior may allow for the euplastic regenerative process to proceed more efficiently, improving long term health of the respiratory system, as evidenced by recent work demonstrating that deletion of the core basal-cell transcription factor Trp63 prevents Krt5+ dysplasia and improves lung function(*8*). Future studies in this direction are warranted in addition to directly promoting the euplastic regenerative response such as recently demonstrated for Wnt signaling(*24*).

Our studies highlight the importance of defining the cellular and molecular differences in acute and chronic lung disease. The presence of iTCHs and their niche signaling activity in acutely injured human lungs from COVID-19 patients and after acute bleomycin toxicity indicates a conserved mechanism of dysplastic repair dominance in these disease states. Conversely, the lack of iTCHs and a reversal of signaling information flows, in particular increased WNT and FGF signaling in degenerative emphysematous diseases, suggests that in these disease states the lung has activated pathways known to promote regeneration. However, given the degenerative phenotype in these diseases, increased WNT and FGF activity is either insufficient or is blocked downstream. This finding points to the need for increased focus on why the alveolar niche does not respond to increased WNT and FGF and indicates the importance of other possible mechanisms driving these diseases including destruction of the supportive matrix in the alveolus and epigenetic regulation of transcriptional networks regulating cell survival and proliferation.

## Methods

### Human subjects

The normal human samples used in this study were from de-identified non-transplanted lungs inititally donated for organ transplantation following an established protocol (Prospective Registry of Outcomes in Patients Electing Lung Transplantation (PROPEL), approved by University of Pennsylvania Institutional Review Board) with informed consent in accordance with institutional and NIH procedures. Consent was provided by next of kin or healthcare proxy. Diseased tissue was obtained from subjects enrolled at the University of Pennsylvania as part of the PROPEL (Penn cohort)(*35*). All selected subjects had diseases listed as classified by a multidisciplinary clinical team. For subjects with COPD (both typical and secondary to alpha-1 antitrypsin deficiency), none had evidence of other parenchymal disease, and none were on systemic steroids. The institutional review board of the University of Pennsylvania approved this study, and all patient information was de-identified before use. While this tissue collection protocol does not meet the current NIH definition of human subject research, all institutional procedures required for human subject research at the University of Pennsylvania were followed throughout the reported experiments. The sample age and pertinent deidentified clinical information are listed in Table S1.

### Mouse lines

All mouse experiments were performed under the protocols approved by the guidance of the University of Pennsylvania Institutional Animal Care and Use Committee. Pdgfra^CreERT2^, Pdgfrb^CreERT2^, Acta2^CreERT2^, Krt5^CreERT2^, Pdgfra^H2B-eGFP^, Rosa26^tdTomato^, Ect2^flox^, and Rosa26^dnMAML-GFP^ mouse lines have been previously described(*17*, *36–42*). All experiments were performed on 8-12 week old mice that were maintained on a mixed C57BL/6 and CD1 background. Both male and female mice were used in all conditions.

### Tamoxifen delivery

For Cre recombinase induction, tamoxifen (Millipore Sigma) was dissolved in corn oil and ethanol mixture (90%/10%, v/v) (Millipore Sigma) to produce a stock solution with a concentration of 20mg/mL. Pdgfra^CreERT2^ and Pdgfrb^CreERT2^ mice were given tamoxifen at a dose of 200mg/kg for 3 consecutive days. Acta2^CreERT2^ mice were given tamoxifen at a dose of 100mg/kg for 4 consecutive days at the timepoints indicated in Figure 2. Krt5^CreERT2^ mice were given multiple doses of tamoxifen (250mg/kg) at days 5 and 8 after influenza, as indicated in Figure S6.

### Influenza infection

PR8-GP33 H1N1 influenza virus was provided by Dr. John Wherry. Two weeks after tamoxifen induction, mice were intranasally administered a dose of 1 LD_50_ (determined empirically in our laboratory) diluted in 50 μL sterile saline as previously described(*23*, *43*).

### Bleomycin administration

Two weeks after tamoxifen induction, 3.5U/kg bleomycin (Teva), diluted in PBS (Thermo Fisher Scientific), was intratracheally administered to anesthetized mice. Control mice received sterile PBS intratracheally.

### Histology, IHC, and RNAscope

Mice were euthanized by CO2 inhalation and the lungs were perfused with ice-cold PBS through the right ventricle. The lungs were then inflated with 2% PFA (Thermo Fisher Scientific) at a constant pressure of 25 cm H2O and were fixed overnight at 4 oC. For human tissue, sections of the peripheral parenchyma approximately 1cm^3^ were dissected from the distal lung and placed in 2% PFA or 4% PFA depending on future use (IHC and RNAscope, respectively) overnight at 4 oC. Tissue was then dehydrated in a series of ethanol concentration gradients, paraffin embedded, and 6um thick sections were cut. Hematoxylin and eosin staining was performed as previously described(*16*).

For IHC, the following antibodies were used on paraffin sections: RFP (goat, Origene, cat# AB8181-200), RFP (rabbit, Rockland, cat# 600-401-379), GFP (chicken, Aves, cat# GFP-1020), SMA (goat, Novus Biologicals, cat# NB300-978), SMA (mouse, Sigma, cat# A5528), SMA (rabbit, Abcam, cat# ab5694), Ki67 (mouse, BD Biosciences, cat# 550609), SP-C (rabbit, Millipore-Sigma, cat# AB3786), Keratin5 (rabbit, Abcam, cat# ab52635), Keratin5 (mouse, Sigma, AMAb91549), Notch1-ICD (rabbit, Cell Signaling, cat# 4147), Jag2 (rabbit, LSBio, cat# LS-C100395), CD45 (rat, Novus Biologicals, cat# NB100-77417).

For RNAscope, lung tissue was prepared, fixed, embedded in paraffin, and sectioned as described above. RNAscope was performed using the Fluorescent Multiplex Reagent Kit v2 (ACDBio) according to manufacturer’s instructions. All RNAscope probes were purchased through ACDBio. The following RNAscope probes were used: Tcf21 (mouse, cat# 508661-C3), Notch3 (mouse, cat# 425171-C2), Pdgfrb (mouse, cat# 411381), Keratin5 (mouse, cat# 415041 and 415041-C2), PDGFRA (human, cat# 604481-C3), PDGFRB (human, cat# 548991-C2), TCF21 (human, cat# 473071), NOTCH3 (human, cat# 558991).

### Imaging and image analysis

Fluorescent images were acquired at 20x and 40x using z-stacks on an LSM 710 laser scanning confocal microscope (Zeiss). Cell counting analysis was completed within FIJI. H&E stained lung sections were tile-scanned using a 4x objective on the Eclipse Ni series upright microscope (Nikon). Whole lobe immunofluorescence tile scans in Figure 5f were acquired using a 4x objective on the Nikon Eclipse Ni series upright microscope.

### Tissue clearing, sectioning, and imaging

Pdgfra^H2B-GFP^:Pdgfrb^CreERT2^:Rosa26^LSL-tdTomato^ mice were given a single dose of tamoxifen as described above. One week later, mice were euthanized, and lungs were perfused with PBS, and inflated with 2% PFA as described above. Lungs were fixed overnight at 4oC with gentle agitation. Lungs were then cleared using the EZclear protocol(*44*). Briefly, the next morning, lungs were washed 4x 30 minutes each in PBS then incubated with 50% (v/v) THF (Sigma-Aldrich) in sterile milli-q water for 16 hours. Lungs were then washed 4x 1 hour each in sterile milli-q water with gentle agitation. Lungs were then washed 4x 1-hour each in PBS then once more overnight at room temperature with gentle agitation. Lungs were then immersed in a three-step sucrose gradient (Fisher Scientific) (10%, 20%, and 30%, prepared in 1x PBS). For each step, lungs were incubated in sucrose overnight at 4 oC with gentle agitation. After incubation with 30% sucrose, lungs were embed in OCT medium (Sakura) and snap-frozen on a bed of crushed dry ice, then stored at −80 oC until sectioned. OCT embedded samples were sectioned (200um) on a cryostat (Epredia, HMH525 NX) and then placed in ice-cold PBS overnight. The next day, sections were placed in EZ view sample mounting and imaging solution (80% Nycodenz (Accurate Chemical & Scientific), 7M urea (Sigma-Aldrich), 0.05% sodium azide (Sigma-Aldrich), in 0.02M sodium phosphate buffer) overnight at room temperature. The next day, cleared sections were mounted on a glass slide in EZ view mounting media and imaged using a LSM 710 laser scanning confocal microscope using a 25x oil immersion objective. Images were analyzed in ImageJ.

### H&E assessment program

To assess injury severity based on H&E staining, we used a modified version of a previously published algorithm(*16*). Using the EBImage package with R(*45*), we imported images and converted RGB pixel intensity to a matrix, we then applied K-means clustering to segment out, in an unbiased manner, 4 different regions of injury severity: normal, moderate, severe, and background. These regions where then color coded as follows: blue, green, red, and black. Using the area of the lobe outlined in H&E, the percent of each injured zone (normal, moderate, and severe) where then calculated as the percent of total lung lobe area. This R script has been uploaded to the Morrisey Lab Github page (https://github.com/Morriseylab).

### Lung tissue digestion, flow cytometry, and FACS

Lungs were harvested and digested into single cell suspensions using collagenase I, dispase, and DNase I, as previously described(*11*). Red blood cells were removed with ACK lysis buffer and then the cell suspension was stained with antibodies. The following antibodies were used for flow cytometry and cell sorting: CD45-PeCy7 (ThermoFisher, clone: 30-F11, cat# 25-0451-82), CD31-APC (ThermoFisher, clone: 390, cat# 17-0311-82), and CD326(aka EpCAM)-APC (ThermoFisher, clone: G8.8, cat# 17-5791-82); DAPI was used to gate out dead cells. All cells were sorted using a 100um sized nozzle. All cells were sorted into ice-cold FACS buffer (PBS, 25mM HEPES (Thermo Fisher Scientific), 2mM EDTA (Invitrogen), and 2% Fetal Bovine Serum (FBS) (Deville)). Flow cytometry and cell sorting experiments were performed using the cell analyzer LSR Fortessa (BD Biosciences) and the cell sorter Aurora CS (Cytek).

### scRNA-seq and analysis

Lineage-traced cells were sorted by gating DAPI^-^, CD45-PeCy7^-^, CD31/326-APC^-^, tdTomato^+^ populations. The tdTomato+ cells were then loaded onto the 10x Chromium controller (10X Genomics). Epithelial and endothelial cells were sorted by gating DAPI-, CD45-PeCy7^-^, CD31/326-APC^+^. Immune cells were sorted by gating DAPI-, CD45-PeCy7^+^. Immune cells were then spiked back in with the epithelial and endothelial cells at a concentration of 25%. The resultant epithelial, endothelial, and immune cell mixture was then loaded onto the 10x Chromium controller (10x Genomics).

For the whole-lung scRNAseq presented in Figure 6, mouse lungs were digested and single cell suspensions were prepared as described above. Cell suspensions were then incubated with magnetic CD45 microbeads (Miltenyi Biotec, 130-052-301) for 30 minutes on ice. Cells were then passed through a magnetic column (LS column, Miltenyi Biotech) to deplete the cell suspension of CD45+ immune cells. CD45+ immune cells were then later recovered and spiked back into the CD45-depleted cell suspension at a concentration of 20%.

For human scRNAseq experiments, lungs were processed and CD45 cells were depleted using the MACS LS collumns as previously described(*46*). CD45 positive cells were not spiked back for human samples. The resultant cell mixture as then loaded onto the 10x Chromium controller.

All samples were loaded to aim for a recovery of 10,000 cells, and libraries were prepped according to the manufacturer’s protocol using the Chromium Single Cell 3’ v3.1 chemistry. Libraries were then sequenced on an Illumina Novaseq 6000 instrument. The sequenced data was processed by aligning reads and obtaining unique molecular identifiers (UMIs) using STARsolo (v2.7.9a)(*47*). For scRNA-seq from lineage-traced samples, libraries were aligned to a modified mm39 mouse genome to include the tdTomato reference sequence. The scRNA-seq data was further processed and analyzed using the Seurat v4 package(*48*). Cells which had less than 200, or greater than 5000, detected genes were removed from the analysis. Cells were also removed if the percent mitochondrial reads were greater than 10%. For the human bleomycin lung single cell data in Figure 7, a percent mitochondrial read cutoff of 30% was used. Feature (gene) data was scaled in order to removed unwanted sources of variation using the Seurat SCTransform function based on percent mitochondrial reads, and the number of genes, and total reads. Integrated libraries were integrated using the RPCA method within Seurat. Non-linear dimension reduction was performed using uniform manifold approximation and projections (UMAPs) (reduction = ‘pca’ and n.neighbors = 15) and Louvain graph-based clustering algorithms. Previously published COPD and COVID-19 scRNA-seq data were downloaded from GSE168191 and GSE159585(*46*, *49*).

Marker genes for each celltype were identified using the FindAllMarkers command using the RNA assay within Seurat. Mesenchymal cells were annotated based on their anatomical location in the lung (i.e. alveolar fibroblast 1 and 2). Peribronchial, adventitial, and vascular smooth muscle cells were annotated using previously published canonical marker genes(*10*). Epithelial, endothelial, and immune cells were annotated using marker genes and LungMAP labels(*13*).

Correlation analysis was performed in R. Slingshot trajectory analysis was performed using the SCP package within R(*50*) using default settings anchoring the trajectory at the cluster enriched for AF1 cells from sham lungs. Transcription factor activity scores were assigned to cells using DoRothEA database within R(*18*). GSEA analysis for Notch signaling (GO:0007219) was performed using the SCP package within R using default settings. Ligand-receptor analysis was performed using CellChat v1(*20*). Prior to ligand-receptor analysis, each cell type was randomly down sampled to 300 cells per cell type to minimize artifacts arising due to discrepancy in cell numbers. After downsampling, CellChat was performed with default settings. We performed ligand-receptor analysis using the ‘Secreted Signaling’ and ‘Cell-Cell Contact’ ligand-receptor databases, excluding ‘ECM-Receptor’ signaling. For the Notch signaling network in Figure 7, both immune and endothelial cells were removed from the ligand-receptor analysis to allow focus on mesenchymal-epithelial niche signaling in disease. Differential ligand-receptor analysis in Figure 6 was performed using CellChat with default settings. GO and KEGG pathway analysis was performed using the clusterProfiler R package using default settings(*51*). WNT ligand module scores in Figure 7 were generated using the AddModuleScore function within Seurat using all human canonical WNT ligands. Normalized module scores were then converted into a heatmap using the heatmap function within base R. Visualization of corrplot, Slingshot, transcription factory activity scores, GSEA, and CellChat were generated using functions within each software package, base R, or ggplot2. UMAPs, violin plots, and dot plots were generated using both Seurat and scCustomize.

### micro-computed tomography (uCT) and analysis

uCT images were generated using a X-Cube CT scanner (Molecubes) located at the Small Animal Imaging Facility at Penn Medicine. Live mice were anesthetized with isoflurane and images were acquired using a respiratory gating approach. The settings for image acquisitation were as follows: reconstructed voxel size: 100 um, X-Ray tube voltage: 50 kVp, X-Ray tube current: 440 uA, exposure time: 32 ms, type of filter used: 0.8 mm aluminum filter, degree of rotation stepping: 0.75 degrees (480 total steps in a 360 degree scan). Reconstructed uCT images were analyzed within Horos.

### Culture of primary post-influenza basal cells

Primary basal cells were isolated from the distal lung of Krt5^CreERT2/+^; Rosa26^dnMAML-GFP/+^ mice at day 22 post-influenza (no tamoxifen was administered prior to injury), using a previously described lung digestion protocol(*8*). Cells were initially plated on 10% Matrigel coated plates and expanded in PneumaCult™-Ex Plus Medium (Stemcell Technologies) + 1x PneumaCult™-Ex Plus Supplement (Stemcell Technologies) + 1:1000 hydrocortisone stock solution (Stemcell Technologies) + 1μM A8301 (TGFb inhibitor, Millipore Sigma) + 10μM Y-27632 (ROCK inhibitor, Cayman Chemical) + 1:5000 Primocin® (Invivogen), which specifically facilitates the outgrowth of basal cells. Cells were treated with DBZ (HY-13526, MedChemExpress) or 4-hydroxytamoxifen (3347905, Cayman) as described in Figure S6.

### Post-influenza Basal Cells Monolayer Colony formation

Cells isolated as described above were seeded at 500 cells per well in growth medium in 6-well plates coated with 10% Matrigel in PBS. The medium was changed to the indicated growth medium supplemented with DMSO, 500 nm 4-hydroxytamoxifen (4OHT), 10 µm DBZ, or 20 µm DBZ after 24 hours. Cells were fixed 6 days later in 3.2% PFA and washed twice with PBS, followed by staining with crystal violet (0.5% crystal violet in 20% methanol) for approximately 10 min to visualize colony formation. The number of colonies formed were counted within FIJI (ImageJ).

### Post-influenza Basal Cells EdU Proliferation analysis

Cells were seeded at a density of 5000 cell/mL on glass cover slips coated with 10% Matrigel in PBS in a 24-well plate. Cell culture media was changed to the indicated growth medium with the DMSO, 500 nm 4OHT, 2 µm DBZ, 5 µm DBZ, 10 µm DBZ 24 hours after plating. After 48 hours, the Click-iT™ EdU Imaging Kit (Thermo-Scientific, C10086) was used according to manufacturer’s protocol to detect the EdU+ cell numbers and the DAPI cell number. Images were acquired using LAS X software with a 20x objective (Leica). The cell numbers were counted within FIJI (ImageJ).

### Post-influenza Basal Cells Organoid Size

Cells were seeded inside 200 µL Matrigel on a 48 well plate at a density of 50 cells/uL. Pneumacult media with supplements was added after the Matrigel solidified at 37 oC. 24 hours after plating, fresh media containing DMSO, 500 nm 4OHT, or 10 µm DBZ was added. Brightfield images were acquired using LAS X software with a 10x objective. Organoid diameters were measured at day 7 using FIJI (ImageJ).

### RNA isolation, cDNA synthesis, and qRT-PCR

For qRT-PCR, RNA was isolated using the Direct-zol RNA Miniprep Plus Kit (Zymo). cDNA synthesis was performed using iScript reverse transcriptase supermix (Biorad). Gene expression was calculated relative to *Rpl19* (“*L19*”) within that sample and expressed as fold change over the average expression. qPCR was run on an Applied Biosystems QuantStudio 6 Real-Time PCR System (Thermo Fisher Scientific) with PowerUp SYBR Green Master Mix (Applied Biosystems). All primer sets are as listed in Table S2.

## Statistical analysis

An unpaired two-tailed t test was used to compare two groups and a one-way ANOVA with Tukey’s adjustment for multiple comparisons was used when comparing multiple groups. Statistical significance was considered when calculated p-values were less than 0.05. All statistical analyses were completed using Graphpad Prism 9.

## Acknowledgements

We thank the Flow Cytometry Core Laboratory at the Children’s Hospital of Philadelphia and the Cell and Developmental Biology Microscopy Core at the University of Pennsylvania for their technical assistance. This research was supported by the National Institutes of Health (HL164929, HL152194, HL132999, HL148857, HL162683, HL168803 to EEM; HL163398 to MCB; HL164350, HL153539 to AEV; HL155821 to EC) and Longfonds BREATH Consortium (EEM).

## Data and code availability

All newly generated mouse scRNA-seq data have been deposited into GEO (GSE249931). Any additional information required to reanalyze the data reported in this paper is available from the lead author upon request.

---

**Supplemental Figure 1:**
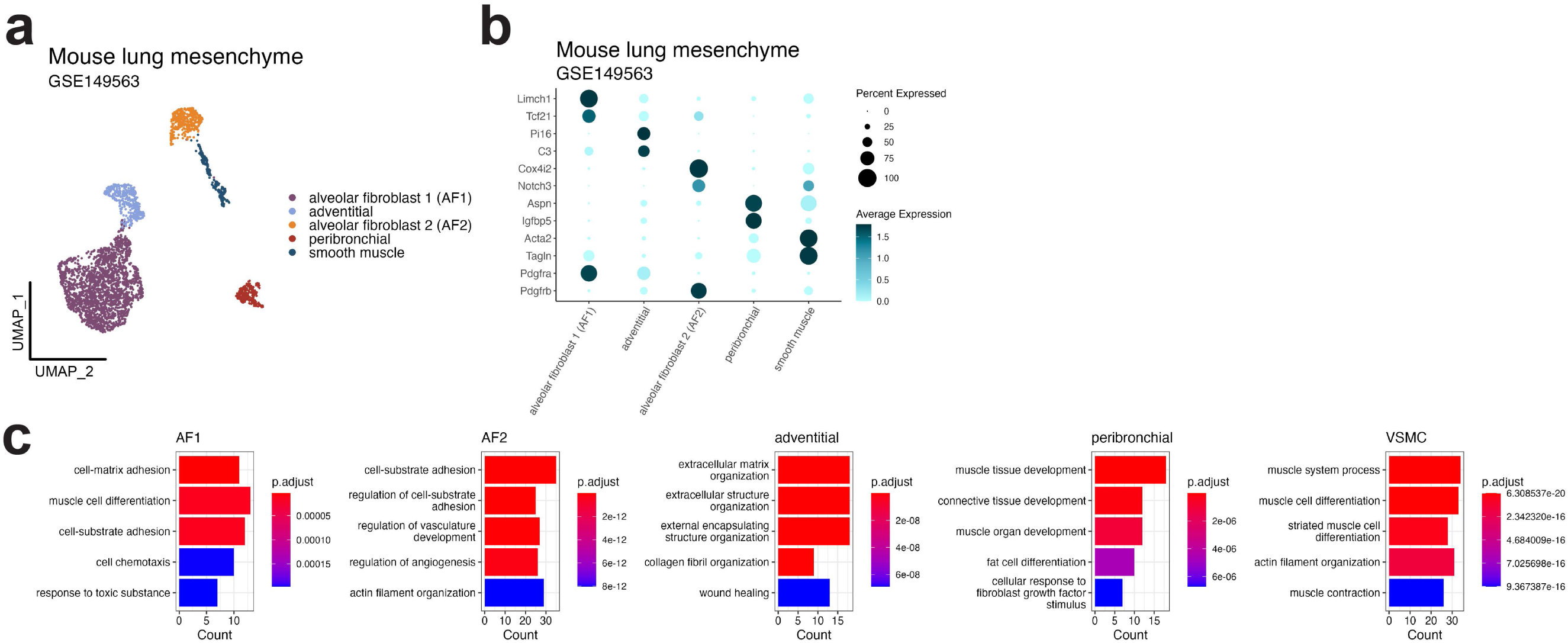
Identification of subtypes of alveolar mesenchyme ie adult mouse lung. (a) UMAP representation of scRNA-seq data from adult mouse lung mesenchyme at homeostasis. Data downloaded from GSE149563(*12*). (b) DotPlot showing relative gene expression of unique marker genes of each mesenchymal subtype in panel (a). (c) GO analysis from the genes uniquely expressed in each cluster in (a). Terms are ranked by adjusted p-values

**Supplemental Figure 2:**
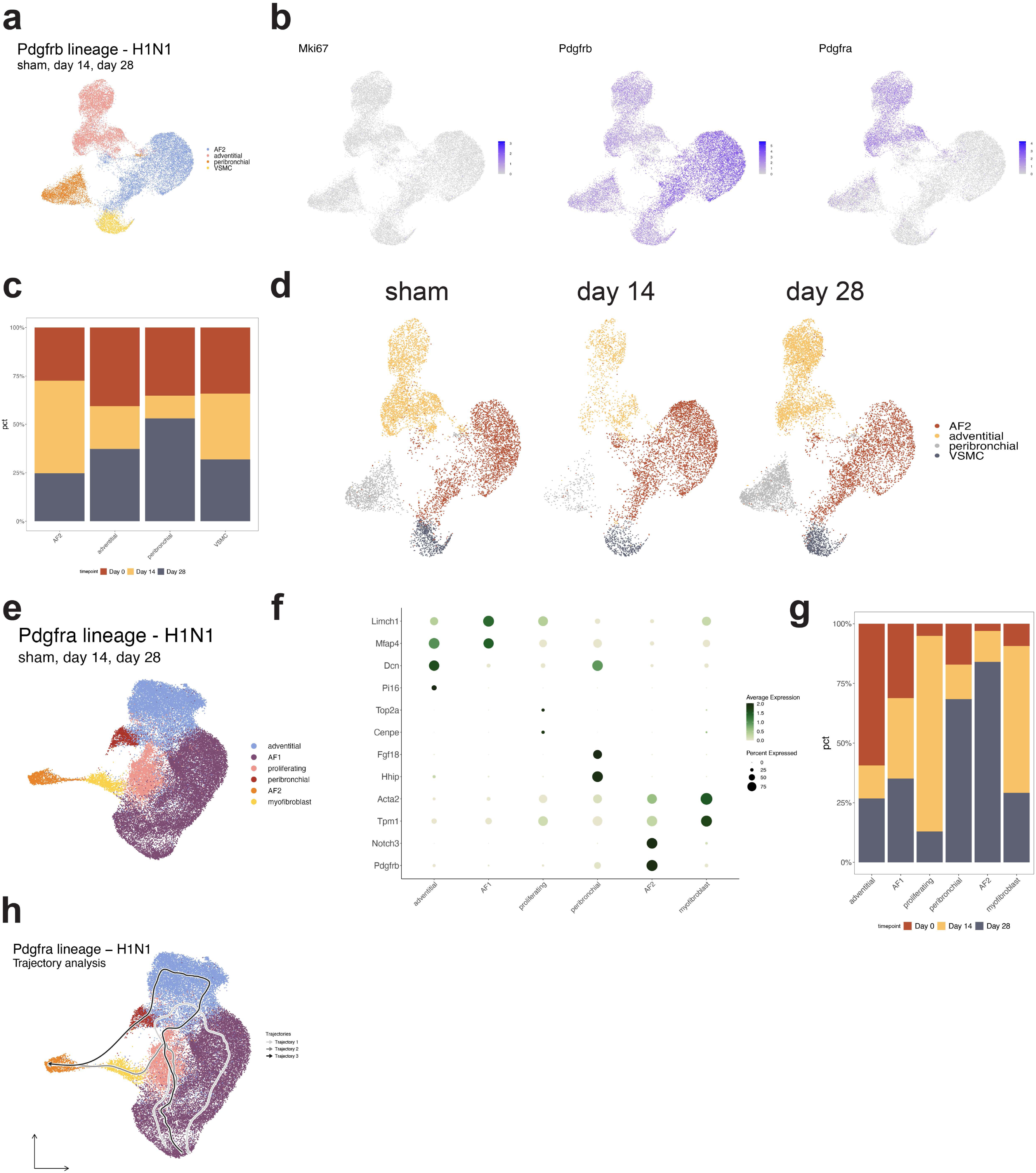
Reactivity of pulmonary mesenchymal cell lineages after respiratory influenza infection. (a) UMAP representation of scRNA-seq data from Pdgfrb+ lineage trace cells after influenza. Data shown represent integratation of sham, day 14, and day 28 timepoints. (b) FeaturePlots showing Pdgfrb+ cells do not proliferate (by Mki67 gene expression) after influenza. Distribution of Pdgfrb and Pdgfra gene expression also shown. (c) Contribution of each timepoint to cell populations shown in panel (a). (d) UMAP separated by timepoint showing that no new cell cluster arises in UMAP space within the Pdgfrb+ cell lineage after influenza infection. (e) UMAP representation of scRNA-seq data from Pdgfra+ lineage trace cells after influenza. Data shown represent integration of sham, day 14, and day 28 timepoints. (f) DotPlot showing relative expression of unique marker genes expressed in cell populations shown in panel (e). (g) Contribution of each timepoint to cell populations shown in panel (e). (h) All Slingshot trajectories generated when AF1s were anchored as starting trajectory point.

**Supplemental Figure 3:**
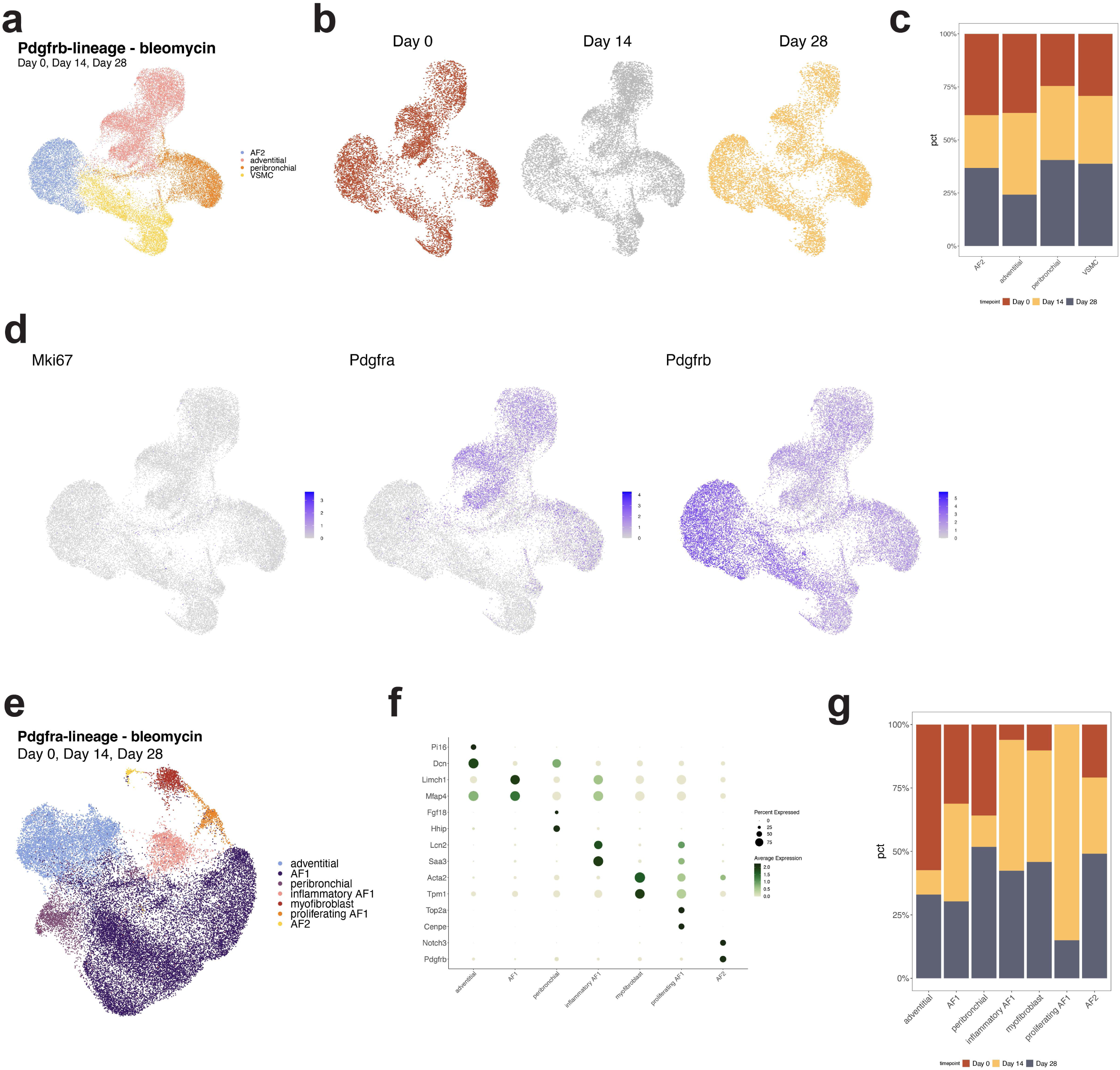
Reactivity of pulmonary mesenchymal cell lineages after bleomycin-induced lung injury. (a) UMAP representation of scRNA-seq data from Pdgfrb+ lineage trace cells after bleomycin. Data shown represent integration of sham, day 14, and day 28 timepoints. (b) UMAP shown in (a) separated by timepoint. (c) Contribution of each timepoint to cell populations shown in panel (a). (d) FeaturePlots showing Pdgfrb+ cells do not proliferate (by Mki67 gene expression) after influenza. Distribution of Pdgfrb and Pdgfra gene expression also shown. (e) UMAP representation of scRNA-seq data from Pdgfra+ lineage trace cells after bleomycin. Data shown represent integration of sham, day 14, and day 28 timepoints. (f) DotPlot showing relative expression of unique marker genes expressed in cell populations shown in panel (e). (g) Contribution of each timepoint to cell populations shown in panel (e).

**Supplemental Figure 4:**
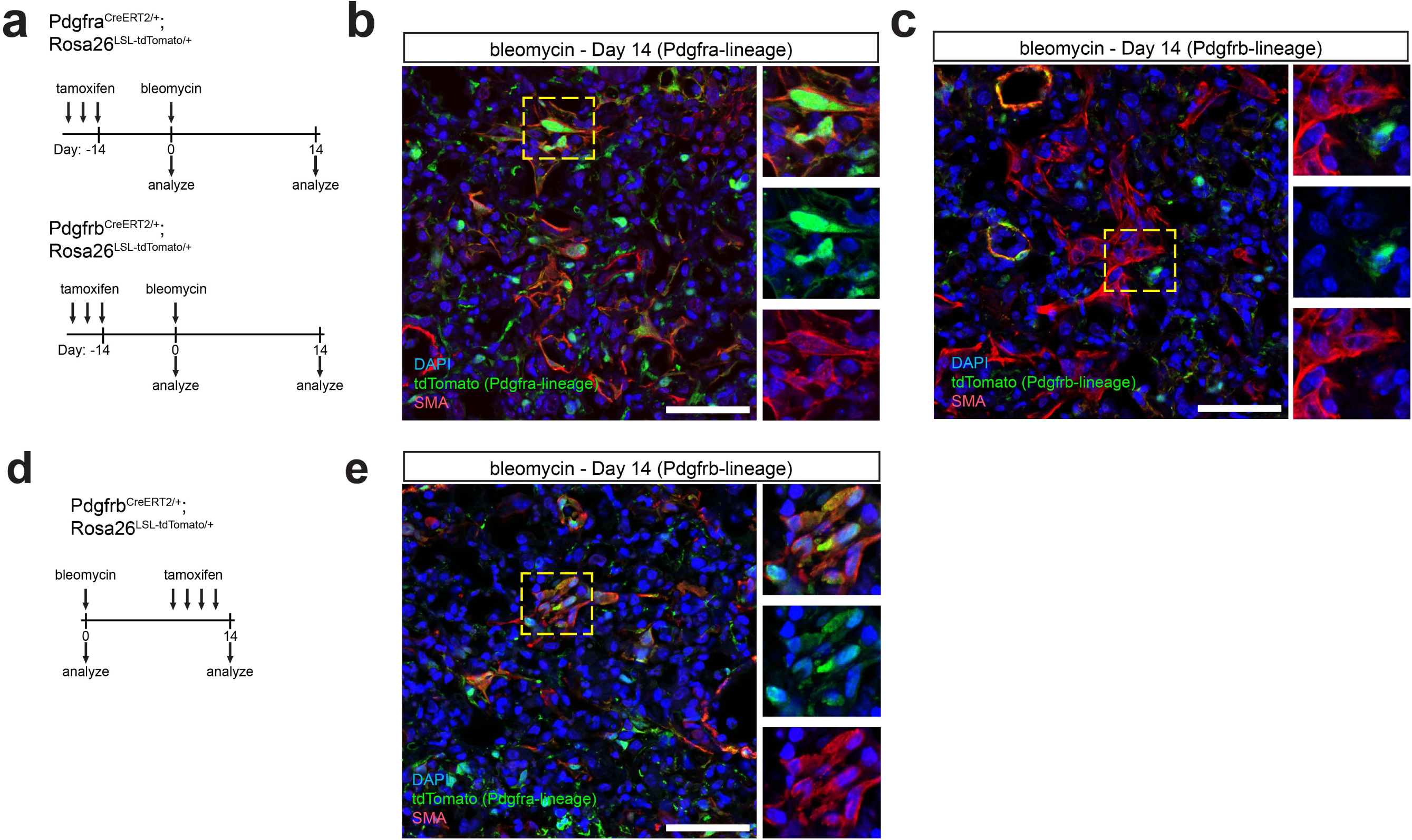
Pdgfra+ cells are primary producers of myofibroblasts after bleomycin-induced lung injury. (a) Experimental schematic showing approach. Pdgfra and Pdgfrb lineage trace mice were given tamoxifen daily for 3 days. 14 days later, mice were given bleomycin (3.5U/kg) and analyzed 14 days later. (b) IHC for tdTomato (lineage-trace) and SMA showing Pdgfra+ lineage trace cells become SMA positive after bleomycin-induced lung injury. (c) IHC for tdTomato (lineage-trace) and SMA showing Pdgfrb+ lineage trace cells do not become SMA positive after bleomycin-induced lung injury. (d) Experimental schematic showing approach. Pdgfrb lineage trace mice were given bleomycin and tamoxifen was delivered 4 days during injury (daily, days 10-13) and mice were collected 1 day later at day 14. (e) IHC for tdTomato (lineage-trace) and SMA showing labeling Pdgfrb+ cells after bleomycin-induced lung injury captures the actively expressing SMA myofibroblasts. All scale bars represent 50 um.

**Supplemental Figure 5:**
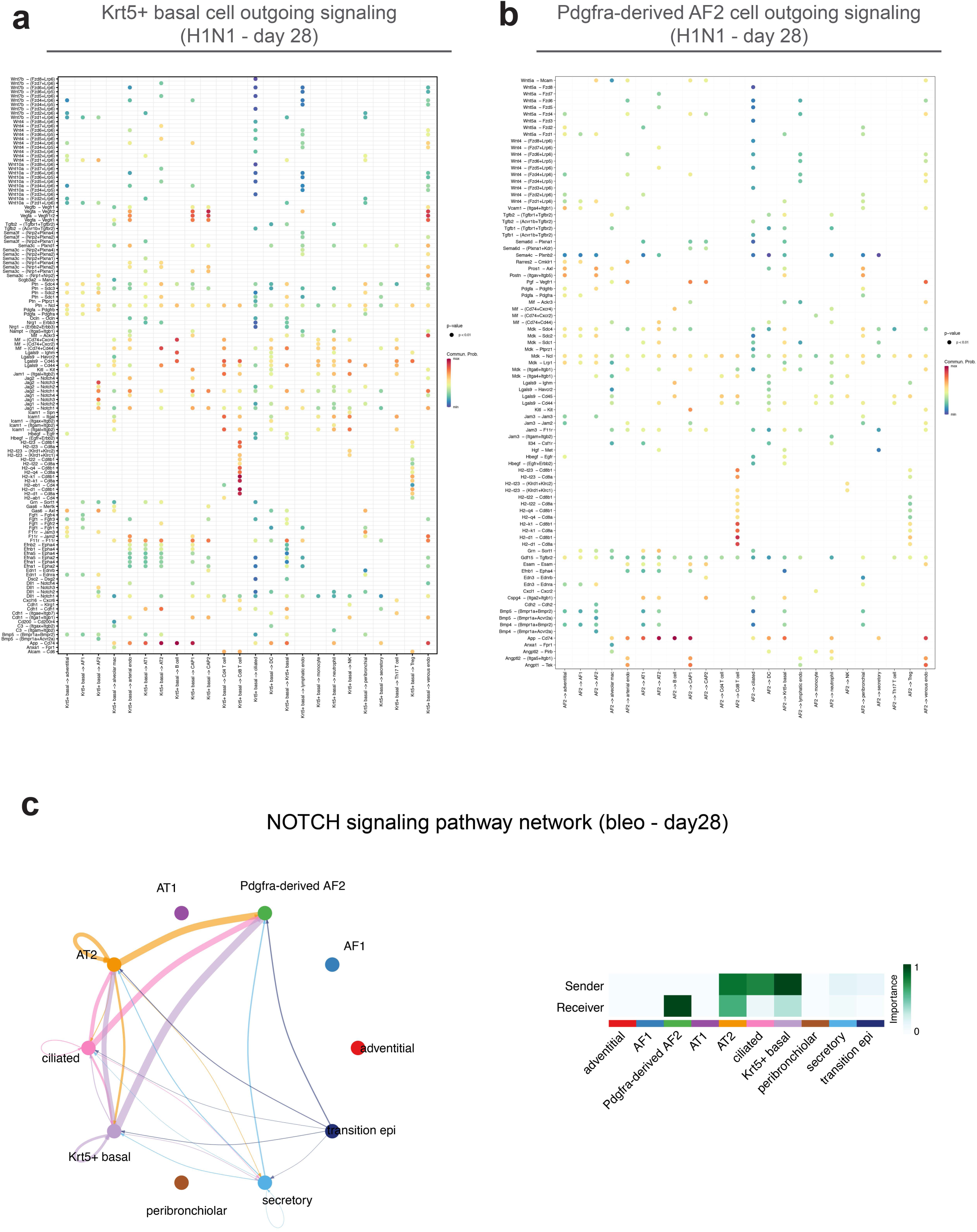
Epithelial and mesenchymal cell crosstalk in the lung after injury. (a) Outgoing cell signaling from the Krt5+ dysplastic basal cells at 28 days after influenza infection showing that Krt5+ epithelium sends the Notch ligands Jag1, Jag2, and Dll1 to the Pdgfra-derived AF2 cells. (b) Outgoing cell signaling from the Pdgfra-derived AF2 cells at 28 days after influenza infection. (c) Notch signaling network between the epithelium and mesenchyme at 28 days after bleomycin-induced lung injury showing Krt5+ dysplastic epithelium and Pdgfra-derived AF2 cells form a Notch niche.

**Supplemental Figure 6:**
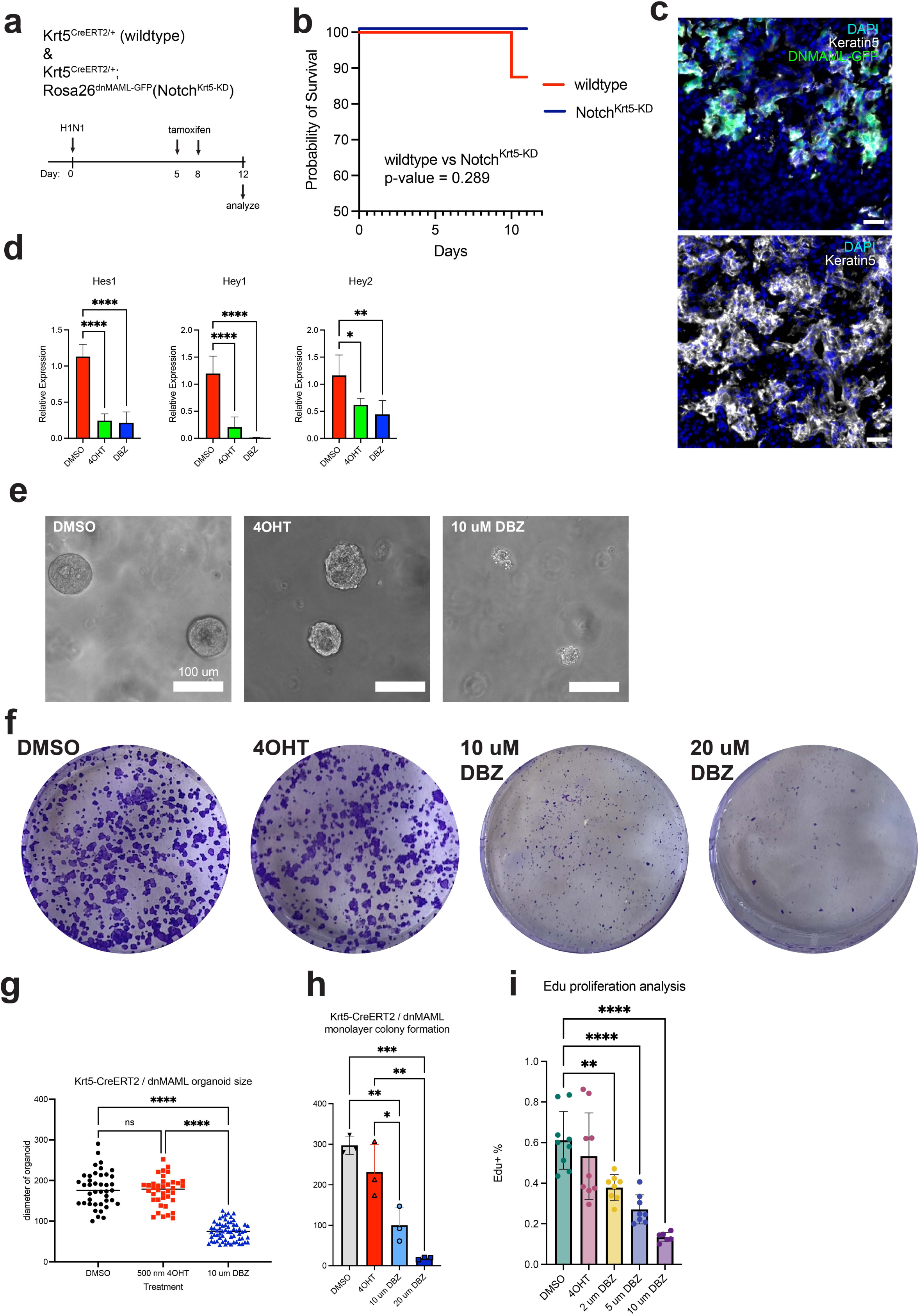
Inhibiting intracellular Notch signaling in the basal cell lineage after flu does not inhibit iTCH formation and basal cell function. (a) Experimental schematic showing approach. Mice were given tamoxifen at days 5 and 8 after influenza and analyzed at day 12. (b) Survival curve showing inhibition of Notch signaling in Krt5+ cells does not improve survival after flu infection. (c) IHC showing Notch^Krt5-KD^ Krt5+ dysplastic epithelial cells migrate and establish an iTCH after influenza. Scale bar represents 20um. (d) qRT-PCR of Notch downstream target genes from cultured in vitro basal cells after exposure to DMSO (vehicle control, n=5), 4-OHT (to genetically activate the dnMAML construct, n=4) or DBZ (small-molecule inhibitor of gamma secretase/Notch signaling, n=5). (e) Representative organoid images of basal cells isolated from Notch^Krt5-KD^ mice after flu showing administration of 4OHT (resulting in intracellular inactivation of Notch) has no effect on organoid size relative to DMSO control. Scale bar represents 100 um. (f) Crystal violet assay showing administration of 4OHT (resulting in intracellular inactivation of Notch) has no effect on colony formation relative to DMSO control. (g) Quantification of organoid sizes shown in panel (d). (h) Quantification of colony formation shown in panel (e). (i) EdU proliferation analysis showing administration of 4OHT (resulting in intracellular inactivation of Notch) has no effect on basal cell proliferation relative to DMSO control. Data points in (f) represent individual organoids, (g) technical replicates in wells. Error bars represent standard deviation. *P<0.05, **P<0.01, ***P<0.001, ****P<0.0001, evaluated by one-way ANOVA with Tukey’s adjustment for multiple comparisons.

**Supplemental Figure 7:**
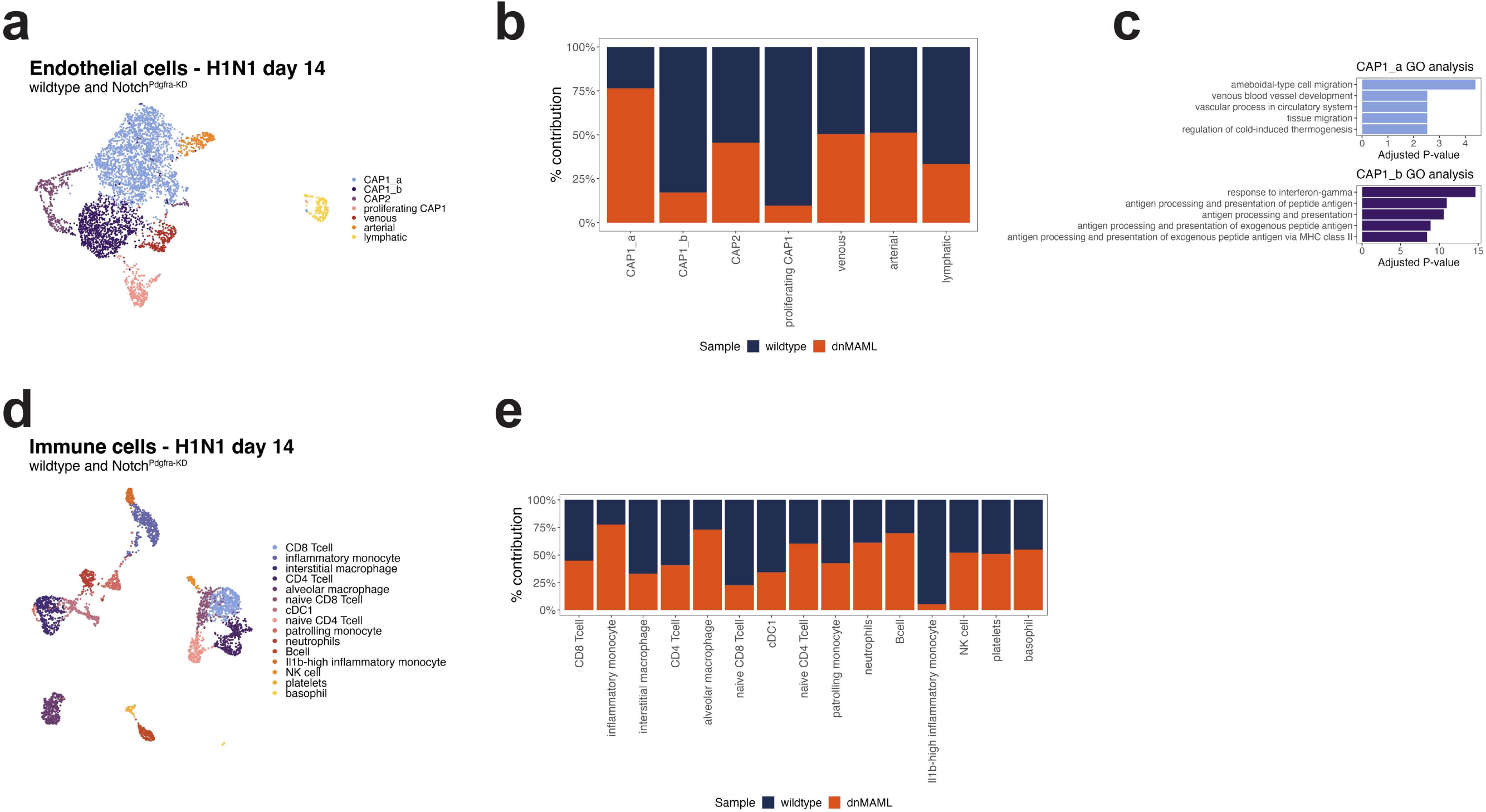
Response of endothelial and immune cells to inactivation of intracellular Notch siganling in Pdgfra+ cells. (a) UMAP representation of scRNA-seq data from endothelial cells at 14 days after influenza infection. Data shown represent merged wildtype and Notch^Pdgfra-KD^. (b) Contribution of genotype/sample to each cell population shown in panel (a). (c) GO analysis from the genes uniquely enriched in CAP1_a and CAP1_b cell populations. (d) UMAP representation of scRNA-seq data from immune cells at 14 days after influenza infection. Data shown represent merged wildtype and Notch^Pdgfra-KD^. (e) Contribution of genotype/sample to each cell population shown in panel (d).

**Supplemental Figure 8:**
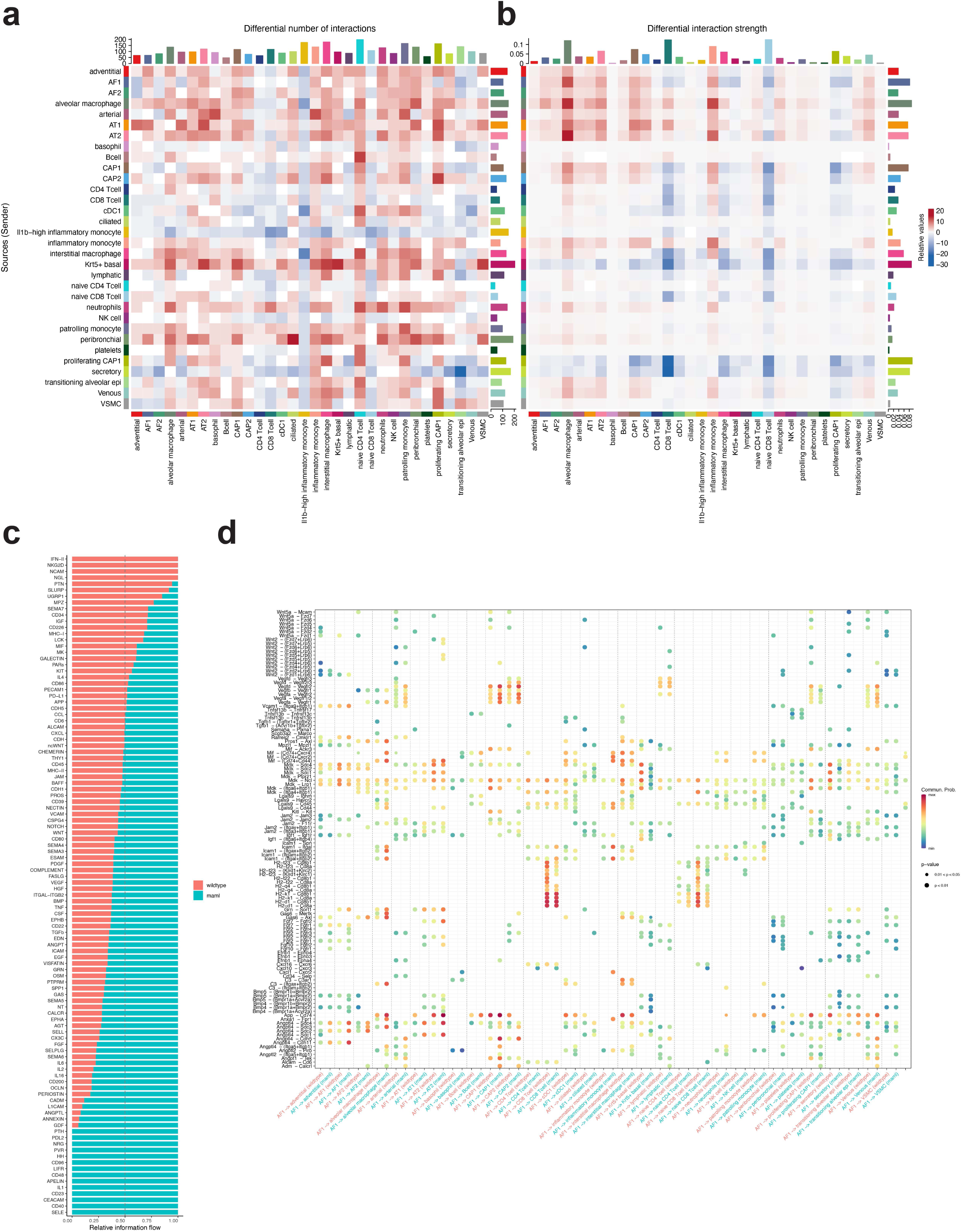
Differential ligand-receptor analysis between wildtype and Notch^Pdgfra-KD^ cell types. (a) Differential number of predicted ligand-receptor interactions between the sending cell (y-axis) and receiving cell (x-axis). Data shown are interaction values relative to wildtype. (b) Differential interaction strength between the sending cell (y-axis) and receiving cell (x-axis). Data shown are interaction strengths relative to wildtype. (c) System wide information flow for the pathways listed. (d) Specific differential ligand-receptor interactions with the AF1 cell as the sending cell. Wildtype = wildtype animals. Maml = Notch^Pdgfra-KD^.

**Supplemental Figure 9:**
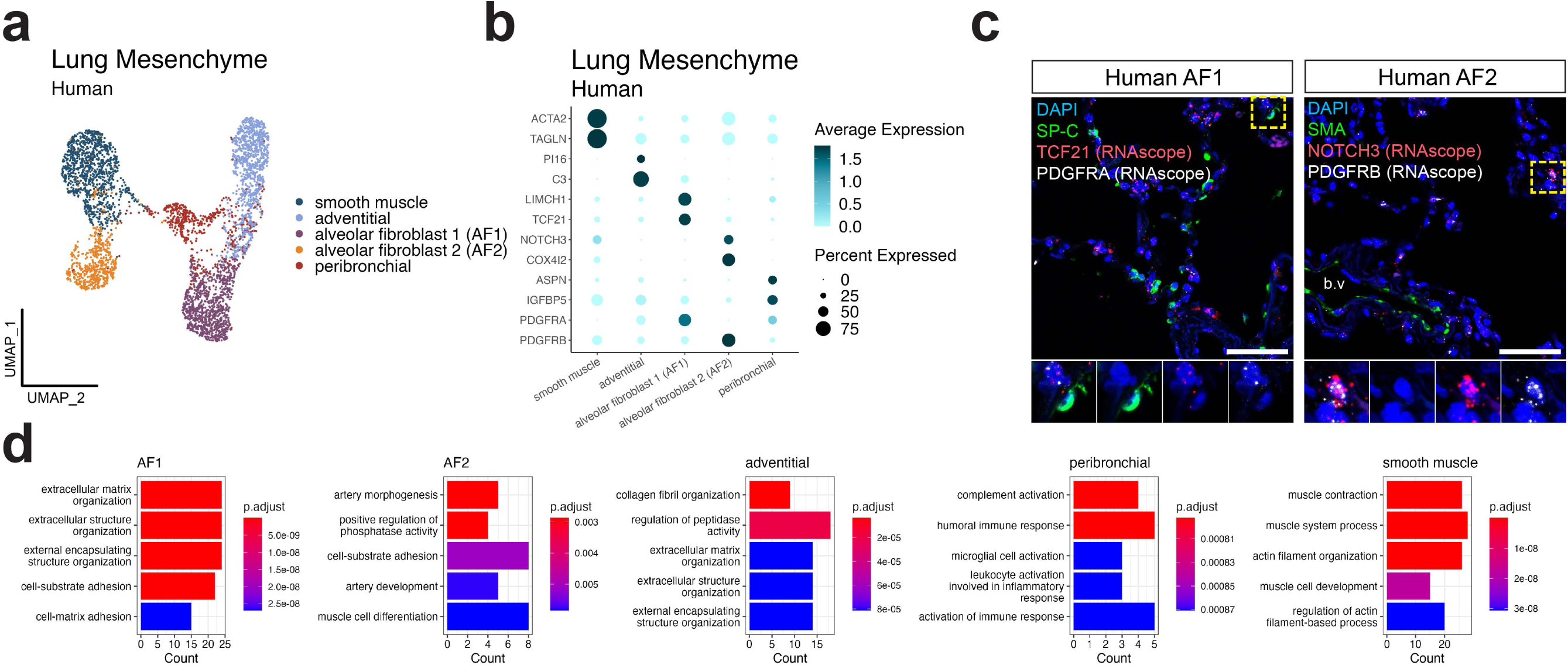
Conservation of alveolar mesenchymal cell types in the human lung. (a) UMAP representation of scRNA-seq data of mesenchymal cells from healthy human lungs (n=13 donors). Data shown represent an integrated dataset of all donors. (b) DotPlot showing relative expression of unique marker genes identified in each cell population in panel (a). (c) IHC and RNA in situ hybridization showing localization of AF1 and AF2 in the human lung. (d) GO analysis from the genes uniquely expressed in each celltype listed in panel (a). GO terms ordered by adjusted p-value.

**<u>Supplemental Table 1: Patient characteristics.</u>**

A list of all of the patients who were included in the analyses presented herein (other than previously published data when mentioned). Age, gender, self-identified race, and cause of death or disease at time of transplantation are provided. Where available, smoking history, FEV1, and/or arterial partial pressure of O2 to fraction of inspired O2 (P/F ratio) is also reported. The use of tissue is indicated by the type of experiment.

**<u>Supplemental Table 2: Primer sequences.</u>**

Primer sequences used to generate the qRT-PCR data in this study.

**<u>Supplemental Data 1: Differential gene expression for Pdgfra-lineage mesenchymal cell types</u>**

**<u>Supplemental Data 2: Differential gene expression for Pdgfrb-lineage mesenchymal cell types</u>**

**<u>Supplemental Data 3: Cleared 200um thick lung section showing separation of Pdgfrb (red) and Pdgfra (green) in the alveolar space.</u>** Each frame is 1 um is in the z-direction. Video is played at 10 frames/second (i.e. 10 um/second).

**<u>Supplemental Data 4: Cleared 200um thick lung section showing separation of Pdgfrb (red) and Pdgfra (green) in the alveolar space.</u>** Each frame is 1 um is in the z-direction. Video is played at 10 frames/second (i.e. 10 um/second).

**<u>Supplemental Data 5: Differential gene expression for integrated scRNA-seq dataset from Pdgfra-lineage trace after influenza infection (sham, day 14, and day 28).</u>**

**<u>Supplemental Data 6: Predicted transcription factor activity within Pdgfra-lineage mesenchymal cell types at day 28 after influenza infection.</u>**

**<u>Supplemental Data 7: Differential gene expression for human mesenchymal cell types.</u>**

