## Supplementary figures and images for "An injury-induced tissue niche shaped by mesenchymal plasticity coordinates the regenerative and disease response in the lung"

### Supplemental Figures

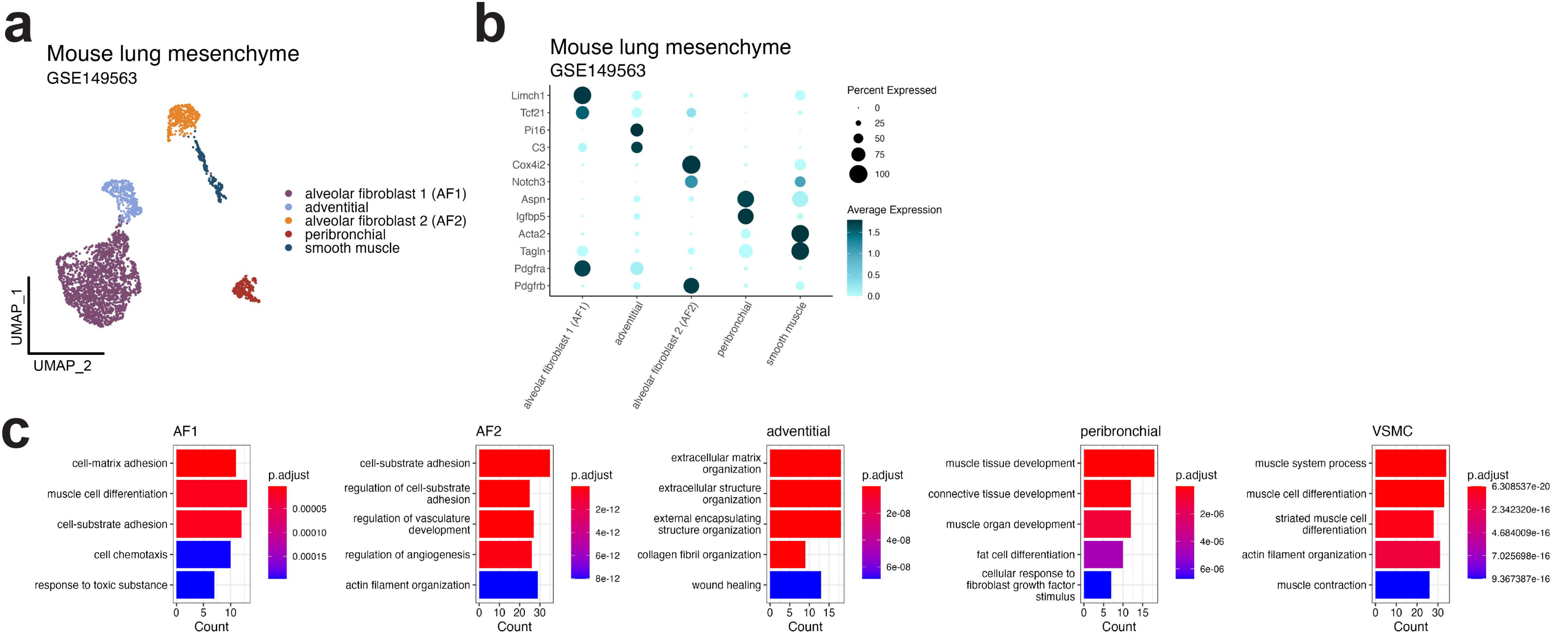

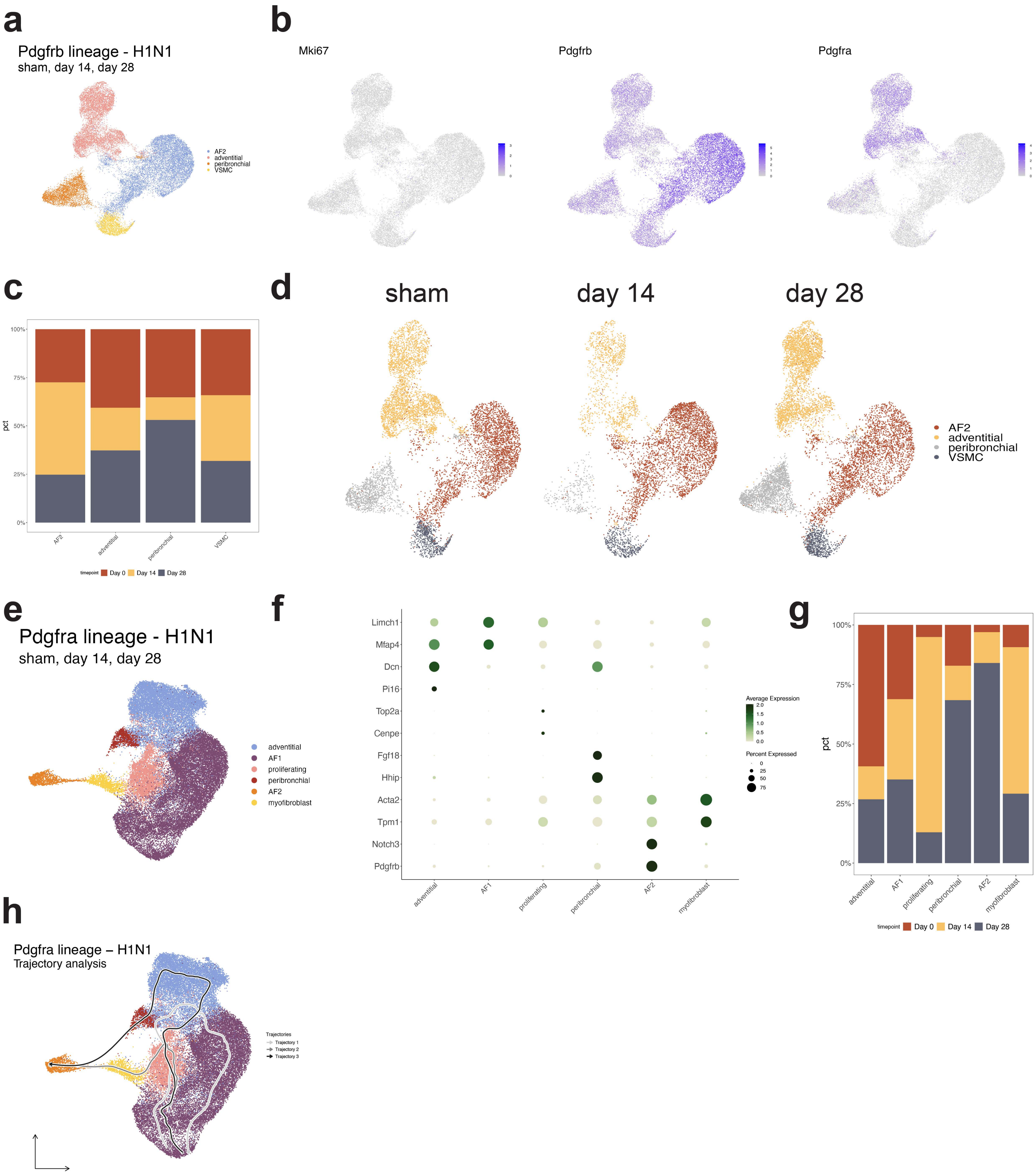

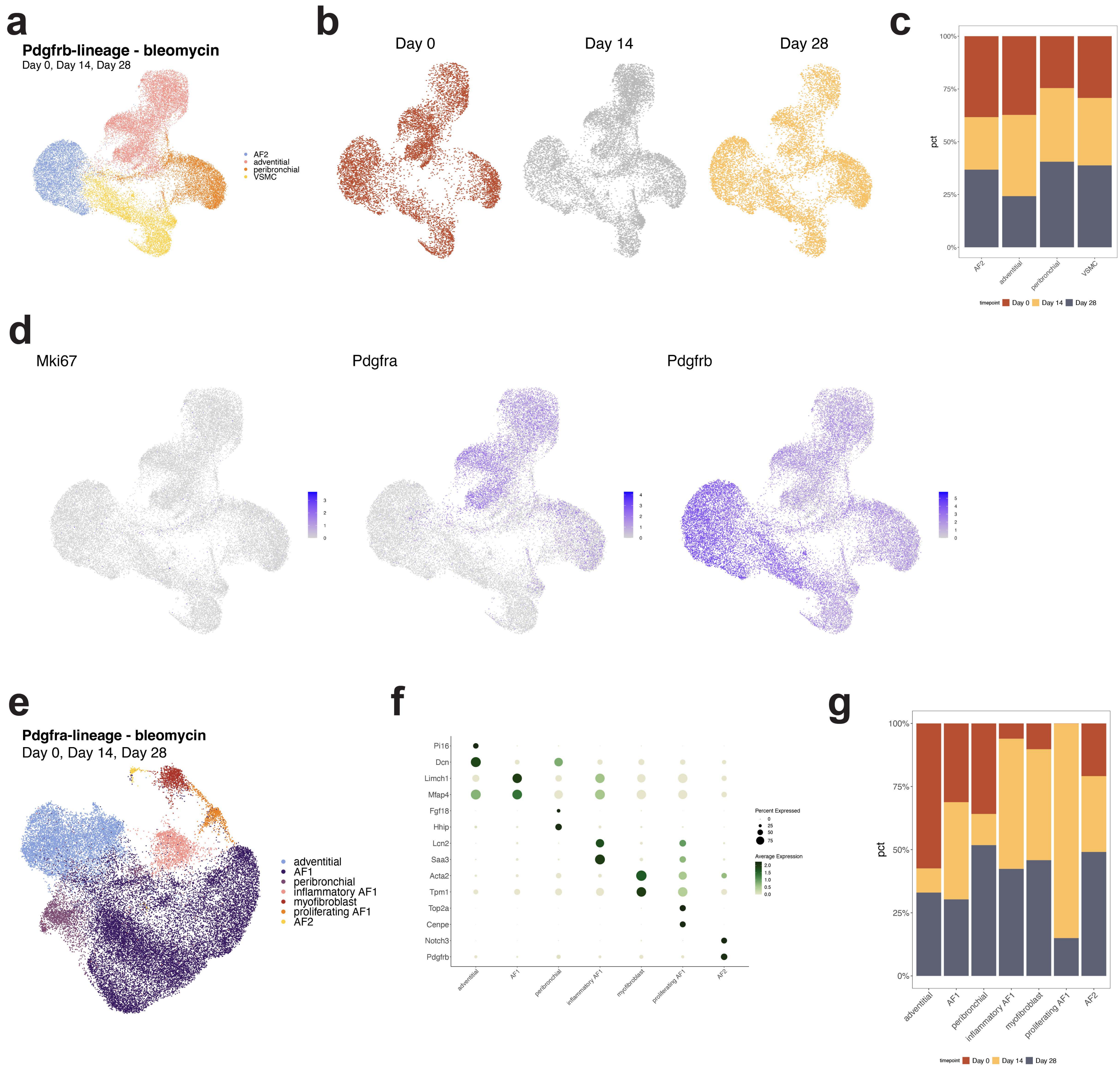

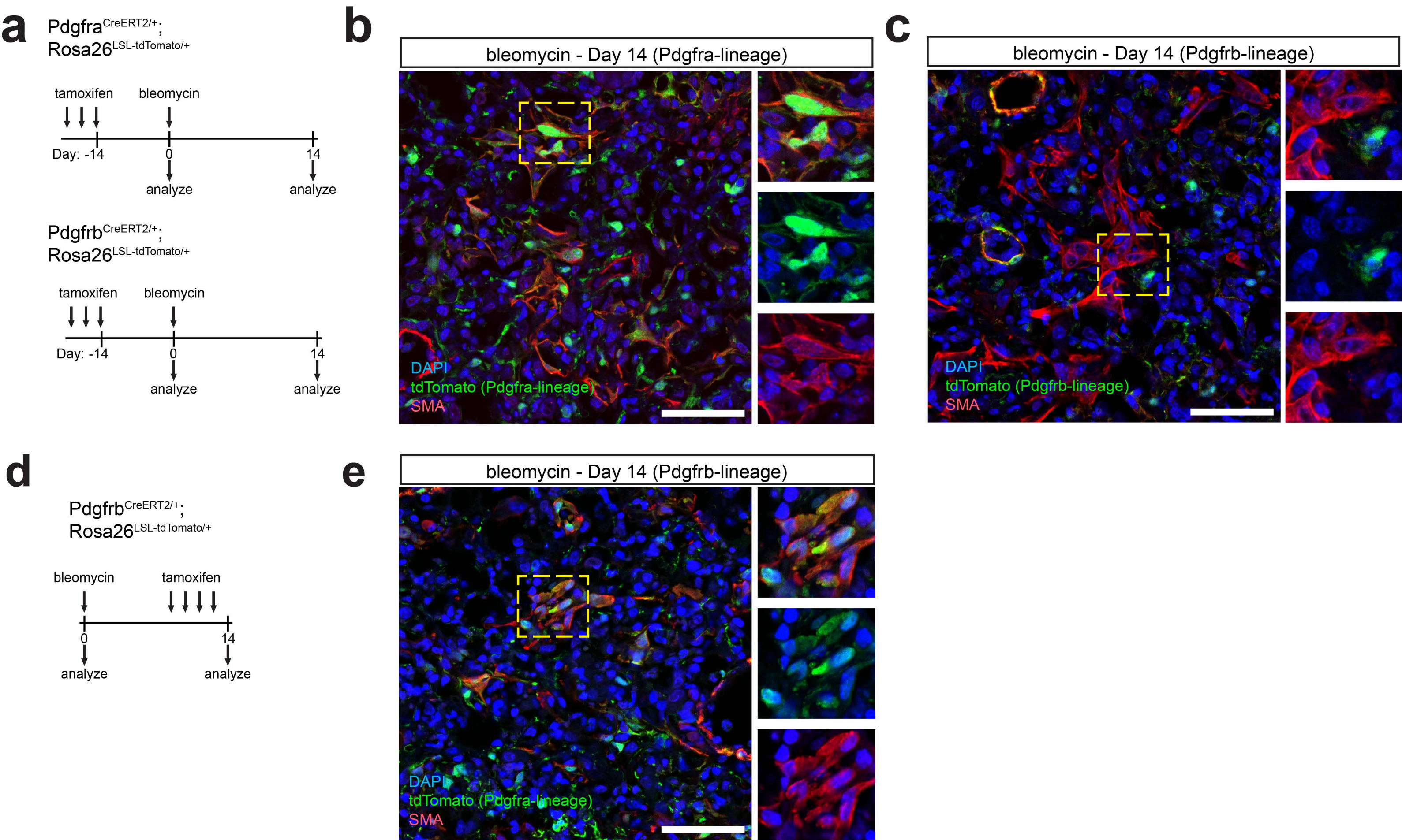

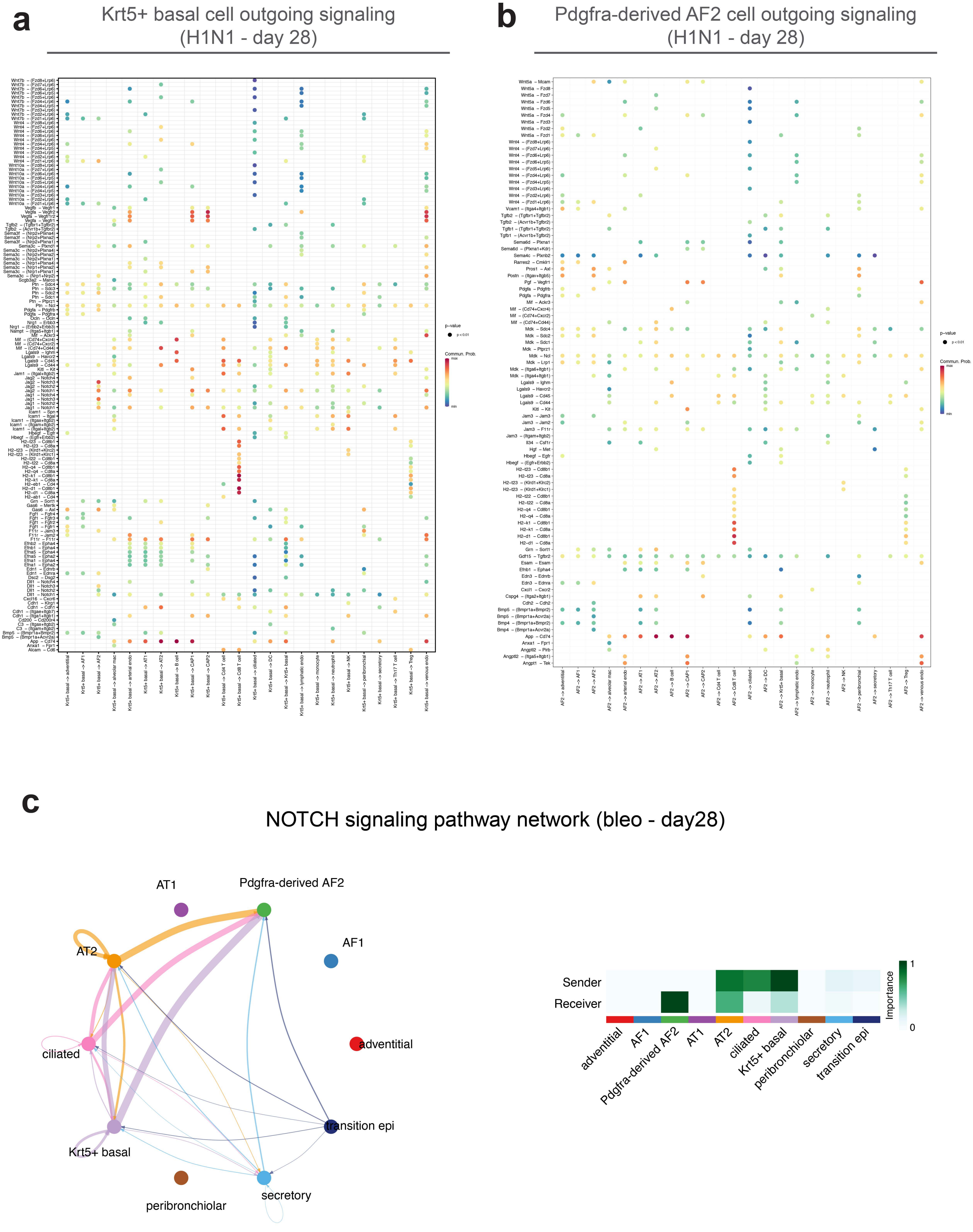

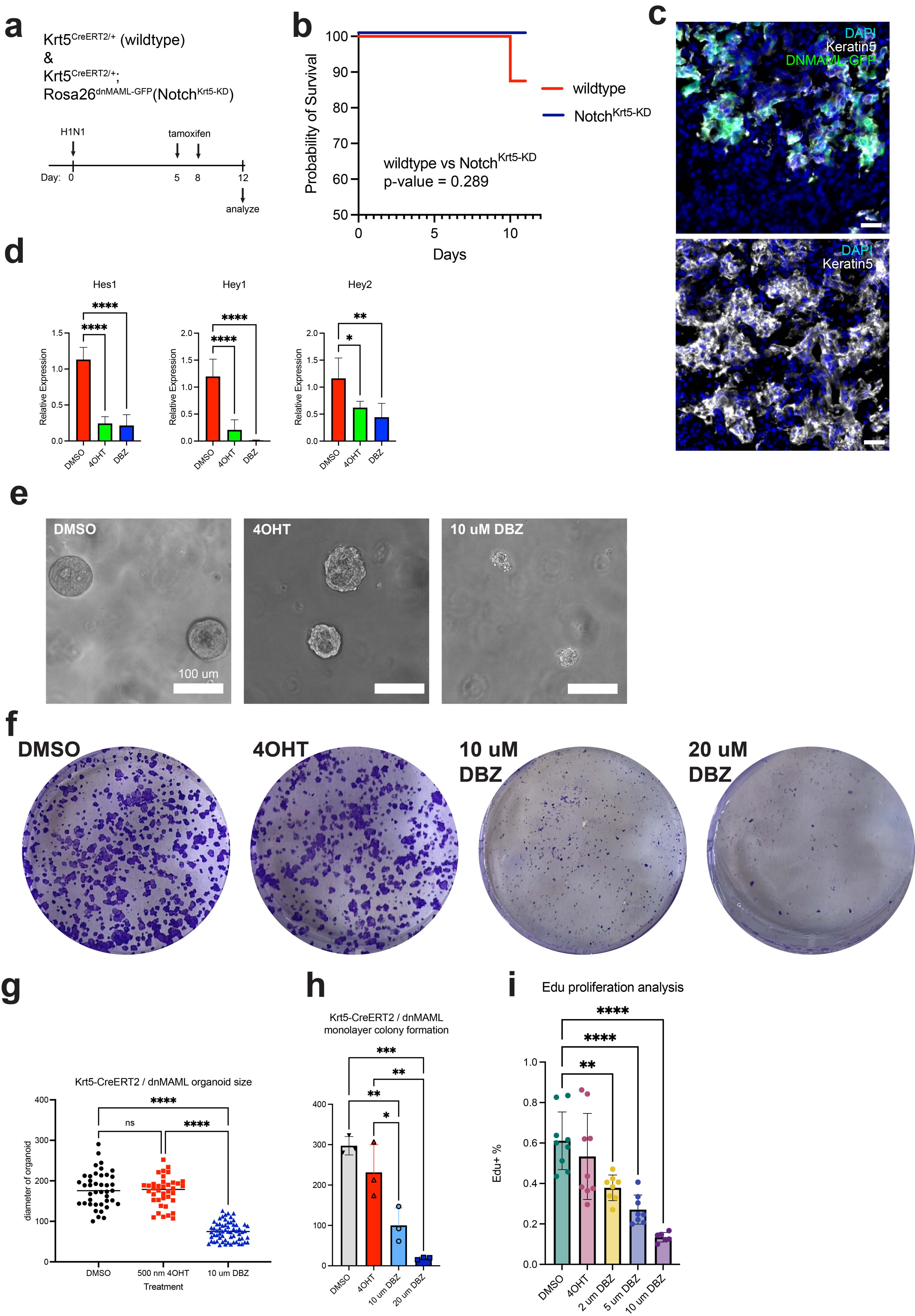

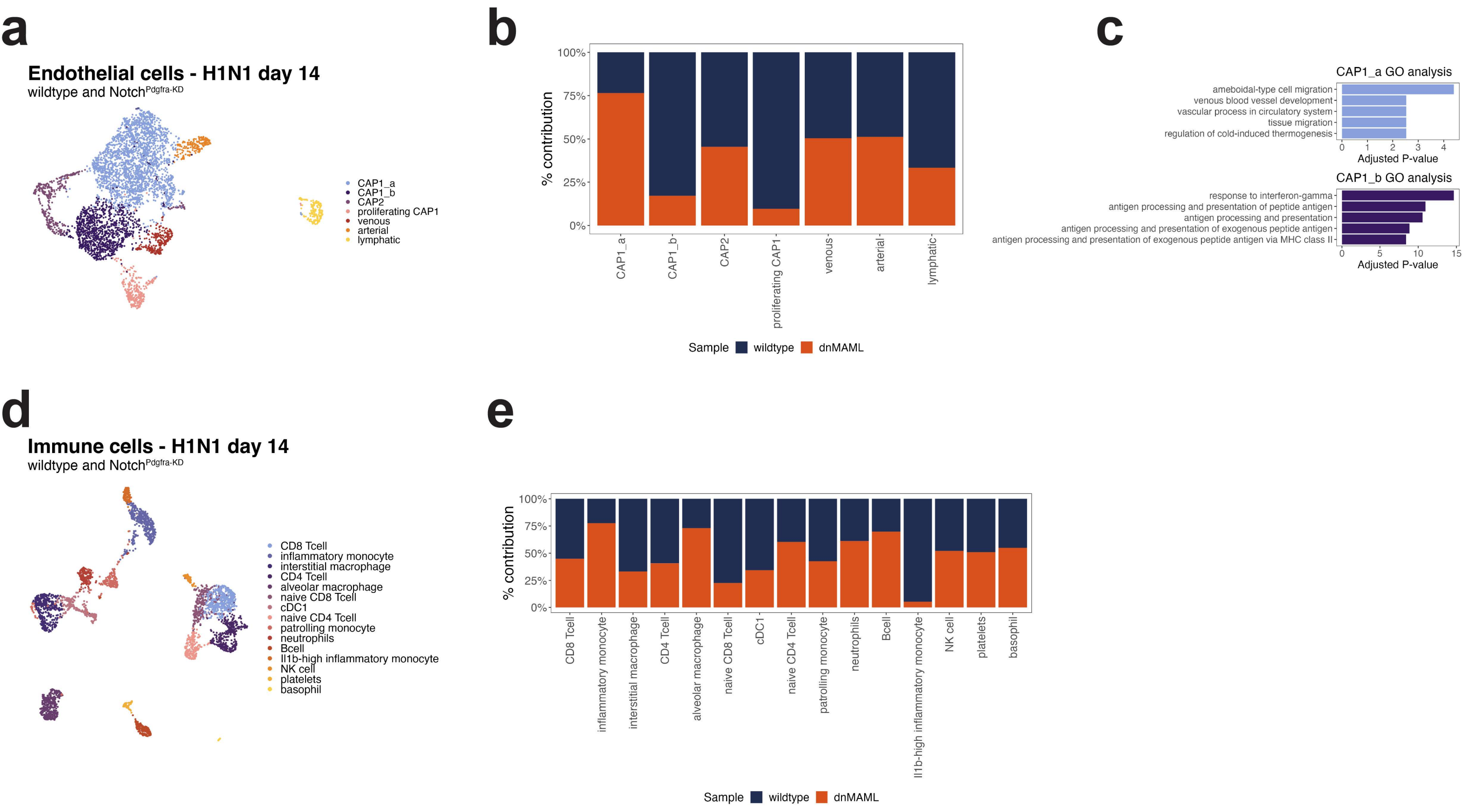

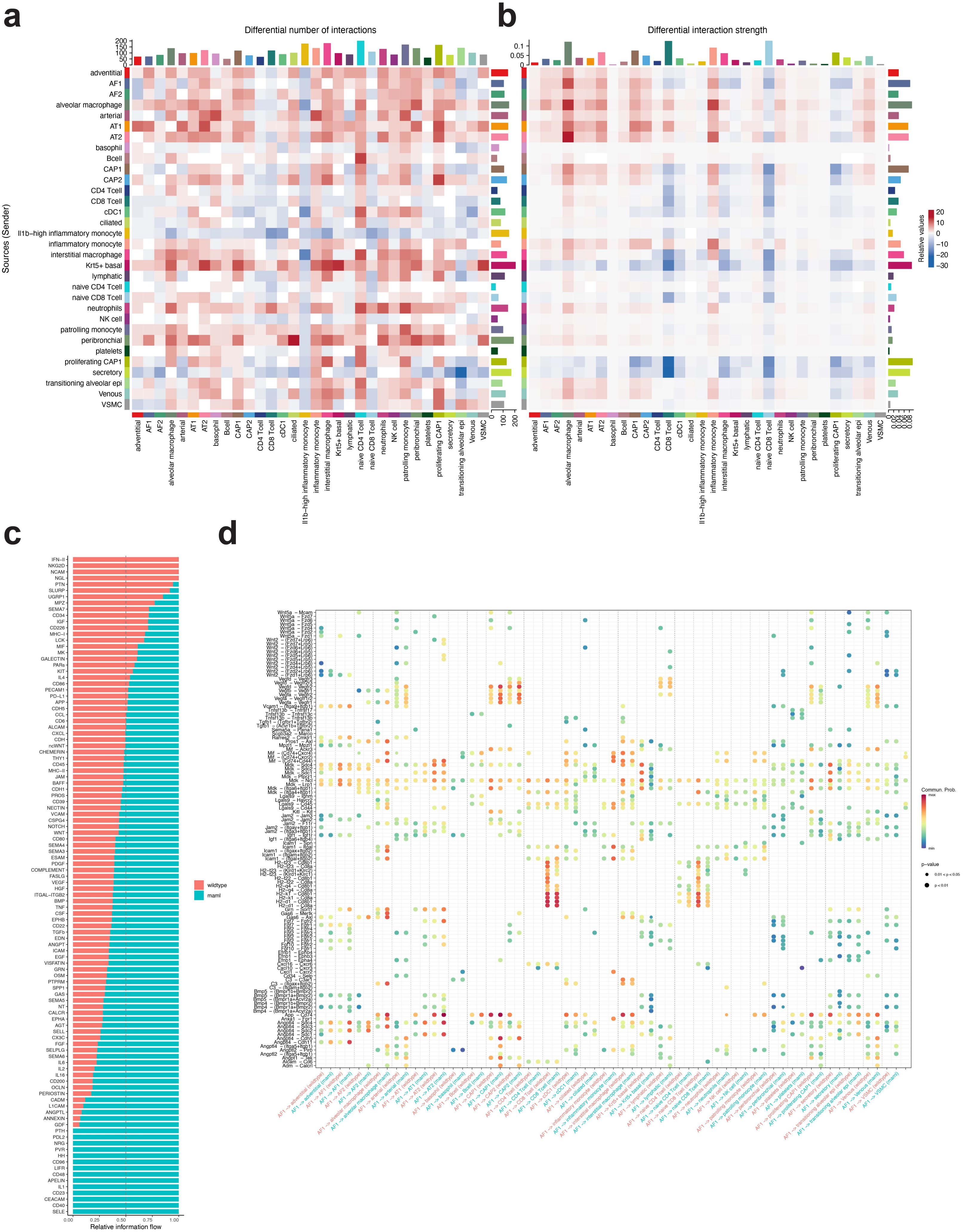

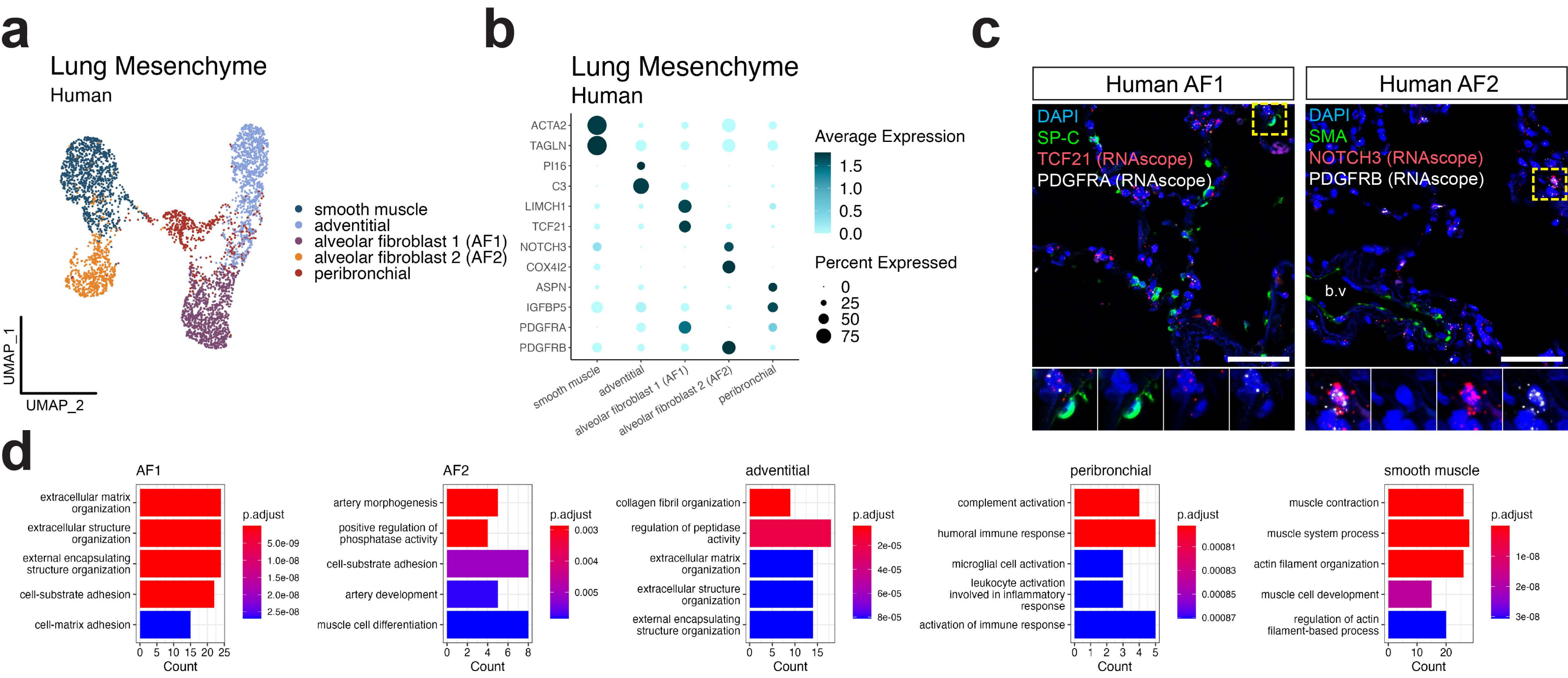
